# Rapid and efficient isolation of intact engineered extracellular vesicles encapsulating functional proteins via internal twin-Strep tetraspanin tagging and Strep technology

**DOI:** 10.1101/2025.11.26.690653

**Authors:** Minh-Tu Pham, Jona Benjamin Krohn, Janis Meyer, Christian Ritter, Martin Schneider, Dominic Helm, Dario Frey, Uta Haselmann-Weiss, Florian Leuschner, Karl Rohr, Ralf Bartenschlager

**Affiliations:** Department of Infectious Diseases, Molecular Virology, Center for Integrative Infectious Diseases Research, Heidelberg University, Heidelberg, Germany; German Center for Infection Research (DZIF), Heidelberg Partner Site, Heidelberg, Germany; Department of Cardiology, Angiology and Pulmonology, University Hospital Heidelberg, Heidelberg, Germany; German Center for Cardiovascular Research (DZHK), partner site Heidelberg/Mannheim, Heidelberg University, Heidelberg, Germany; BioQuant Center, IPMB, Biomedical Computer Vision Group, Heidelberg University, Heidelberg, Germany; Proteomics Core Facility, German Cancer Research Center (DKFZ), Heidelberg, Germany; Division Virus-Associated Carcinogenesis, German Cancer Research Center (DKFZ), Heidelberg, Germany

**Keywords:** affinity-based isolation, intact engineered EV, twin-Strep-tagged CD63/CD9, Strep technology, protein-of-interest, high purity, photoactivatable protein release, virus contamination

## Abstract

Extracellular vesicles (EVs) are important mediators of intercellular communication. Achieving high purity of intact engineered EVs through separation from the overwhelming background of unmodified vesicles and cellular or serum-derived contaminants remains a major challenge, yet is critical for fundamental studies of EV biology and EV-based therapeutics. To address this unmet need, we applied structure-guided engineering to canonical EV tetraspanins. Focusing on CD63 and CD9, we identified sites for internal insertion of the twin-Strep tag via flexible linkers into unstructured regions of their large extracellular loops. This design retains functional properties of the scaffolds while providing a robust molecular handle enabling rapid, efficient, and selective antibody-free affinity-based isolation of intact EVs using advanced Strep technology. Fusing proteins of interest (POIs) to the C-terminus of these scaffolds, either directly or via a photoactivatable protein, allowed POI loading into the lumen of twin-Strep tagged EVs with optional optogenetic control of POI release. Proteomic analysis confirmed the high purity of captured engineered EVs with >96% of contaminants removed, enabling sensitive detection of numerous EV-associated proteins. We also report the efficient removal of contaminating virus particles resembling EVs in size and density. Purified engineered EVs successfully delivered an encapsulated fluorescent reporter into target cells, bypassing dye labeling commonly used to track EVs. We demonstrate endosomal escape of the reporter, facilitated by EV decoration with vesicular stomatitis virus glycoprotein, and cytoplasmic release upon optogenetic activation. Our toolbox may serve as a broadly applicable strategy for the efficient production of intact and highly pure engineered EVs to support potential fundamental or translational EV studies.

## Introduction

Extracellular vesicles (EVs), including exosomes and microvesicles, are lipid bilayer-enclosed nanoparticles secreted by cells and playing pivotal roles in intercellular communication by transferring bioactive molecules, including proteins, nucleic acids, and lipids [1–3]. Their biocompatibility, stability, deep tissue penetration, and ability to deliver cargo proteins make EVs promising platforms for therapeutic delivery of proteins or peptides of interest (POIs). Potential applications comprise regenerative medicine, oncology, immunotherapy, aesthetic medicine, antiviral therapies, and gene editing [4–9].

Despite their potential, the practical use of engineered EVs faces major challenges, particularly in separating engineered EVs carrying the desired cargo from the heterogeneous pool of secreted EVs lacking cargo as well as from contaminants of similar size, density, and molecular composition [3, 10–12]. An even greater concern is the co-purification of EVs with enveloped viruses or virus-derived products present in EV-producing cells [13, 14]. High purity is of particular importance when assigning specific biological functions to EVs, as impurities can substantially confound functional interpretations. In particular, EV corona dynamics and equilibria, defined by the reversible binding and exchange of molecules on the EV surface in biological fluids, especially lipoprotein-EV associations, can obscure EV-specific biological effects [15]. Another major bottleneck for EV applications is the lack of reliable dosing and standardization of engineered EVs, which is largely attributable to impure EV preparations [3, 16]. Furthermore, fundamental mechanistic studies of EV biogenesis, such as the roles of different tetraspanins in EV membrane curvature sensing and cargo sorting, remain challenging because current methods cannot selectively and efficiently enrich intact EV subsets with sufficient integrity and specificity [17].

Conventional isolation methods, including filtration, density gradient centrifugation, size exclusion chromatography, and precipitation methods, are effective for bulk EV recovery but lack the specificity needed to separate engineered EVs from complex particle mixtures. In contrast, affinity-based capture strategies can enrich defined EV subpopulations by exploiting specific molecular interactions. These strategies include antibodies or aptamers targeting EV surface proteins such as CD63, CD81, CD9, or the Tim4 protein, the latter binding phosphatidylserine on the EV surface in a Ca²⁺-dependent manner [18, 19].

Antibody-based EV capture offers relatively high specificity but often yields low recovery, as harsh elution steps compromise EV integrity and bear the risk of co-elution of antibodies. Aptamer-based methods enable gentler elution and better EV preservation, but typically at the cost of lower efficiency [18, 20, 21]. More recently, engineered tagging systems have been explored. The Snorkel-tag system, for example, fuses a transmembrane domain to the EV surface marker protein CD81 along with tags such as HA, Flag, and a proteolytic cleavage site [22]. While effective, this method requires multiple overnight incubations and protease treatment, raising concerns about potential contamination that could affect downstream applications. Similarly, CD63 has been modified with a 3x-Flag tag to enable the pulldown of engineered EVs [23]. Yet, none of the reported methods fully overcome the key challenges: isolating engineered EVs with high yield, specificity, and purity, while minimizing contamination and processing time [18]. Additionally, the removal of excess dyes such as DiI or PKH, which are often used for EV labeling remains a major concern, as residual dye micelles and stained non-EV particles can be taken up by recipient cells [20, 21].

Strep technology, particularly the twin-Strep tag paired with the directed-evolution StrepTactinXT variant, is used to isolate intact protein complexes under native conditions with near-covalent affinity and high purity [24–26]. However, this approach has not been directly applicable to EV scaffolds such as tetraspanins to isolate intact EVs, because tags fused via the N- or C-termini end up in the EV lumen, rendering them inaccessible for capture and preventing the use of Strep-based purification of engineered EVs without detergent-mediated membrane disruption.

Here, we report structure-guided, dual-functional EV scaffolds created by inserting the twin-Strep tag via flexible linkers into an extracellular loop of CD63 and CD9 and fusing POIs to their C-termini. These scaffolds enable the selective Strep-based isolation of engineered EVs while supporting efficient protein cargo loading into the EV lumen and retaining biological activity. We validate this approach by profiling the EV proteome, demonstrating separation from contaminating virus particles and confirm the transfer of functional encapsulated reporter proteins into target cells.

## Results

### Structure-guided insertion of twin-Strep into CD63/CD9 extracellular loops and C-terminal fusion of proteins of interest

To enable affinity-based EV isolation, we engineered CD63 that was selected for several reasons: first, it is a canonical EV-sorting protein consistently expressed across 60 cell lines and represents a core component of EVs from diverse cellular origins [27]; second, when genetically fused to cargo proteins, it ranks among the most effective EV sorting factors [28]; and third, CD63 has been widely used as a benchmark EV marker [17]. We employed the most recent Streptavidin (Strep) affinity technology, i.e. the Strep-TactinXT/twin-Strep-tag system, because it has picomolar ligand – receptor affinity and enables the native isolation of bulky protein complexes with high purity [24–26]. In spite of these superior properties, this technology has not been broadly applied to the internal tagging of proteins, including tretraspanins where C- or N-terminal tags end up in the EV lumen, allowing affinity capture only after EV lysis. To overcome this limitation, we used structure-guided design to identify adequate insertion sites for the twin-Strep (2Strep) tag based on a high-confidence AlphaFold-predicted model of CD63 that closely resembles the solved structures of CD9 and CD81 [29, 30] (Fig. 1A, left panel). We inserted the 2Strep-tag, comprising two tandem copies of the StrepII tag flanked by long flexible linkers, into various positions within the unstructured region of the large extracellular loop (LEL) of CD63 after small amino acid residues (serine residues 159 or 161, or glycine residue 176), or into a site in the small extracellular loop (SEL) after glutamine residue 36, previously reported to tolerate insertions [31] (Fig. 1A, upper right). To enable production and loading of POIs into the lumen of 2Strep-tagged EVs, we fused the POIs to the C-terminus of each of these CD63-2Strep-tag scaffolds (Fig. 1B). For easy read-out, we selected Nanoluciferase (Nluc) (construct #2) or the fluorescent reporter protein mScarlet (construct #5) [32] as POIs (Fig. 1B). The latter was N-terminally fused to a photocleavable protein (PhoCl_2_) [33, 34] and C-terminally to a nuclear localization signal (NLS), respectively. While Nluc enables rapid quantitative measurement of EVs during isolation and uptake into recipient cells, the PhoCl_2_-mScarlet-NLS design allows optogenetic control of mScarlet release from the scaffold via a 405 nm LED light pulse applied to the EV preparation, thereby enabling mScarlet nuclear transport following endosomal uptake and escape after delivery into target cells. As a technical control, we generated a CD63-Nluc fusion construct lacking the 2Strep-tag (construct #1) (Fig. 1B). We generated stable HEK293T and Huh7 cell lines by lentiviral transduction to express the engineered constructs, because both cell types produce CD63-containing EVs and Huh7 cells are particularly well suited for studying endocytic pathways using a range of methods, including imaging-based techniques [2]. To rapidly screen the insertion site best suited for 2Strep-based isolation of EVs, we captured EVs containing the various CD63-2Strep-Nluc proteins from ∼5 mL of cell-conditioned media using 1mL gravity-flow columns packed with Strep-TactinXT resin. The Nluc activity contained in all fractions, including flow-through, washes, and eluates, the latter obtained by using a 50 mM biotin-containing buffer, was measured to assess the binding of the engineered EVs to the resin and the EV yield. The 2Strep-tag insertion at all three positions in the LEL of CD63 allowed robust and comparably efficient CD63 secretion and affinity-based capture (Fig. S1A). While the insertion of the tag into the SEL (Q36-2Strep-Nluc) also allowed efficient secretion, binding to the matrix appeared weak, with most material being lost in the first washing and less than 1% of the input Nluc-activity being retained in the eluates.

**Figure 1.**
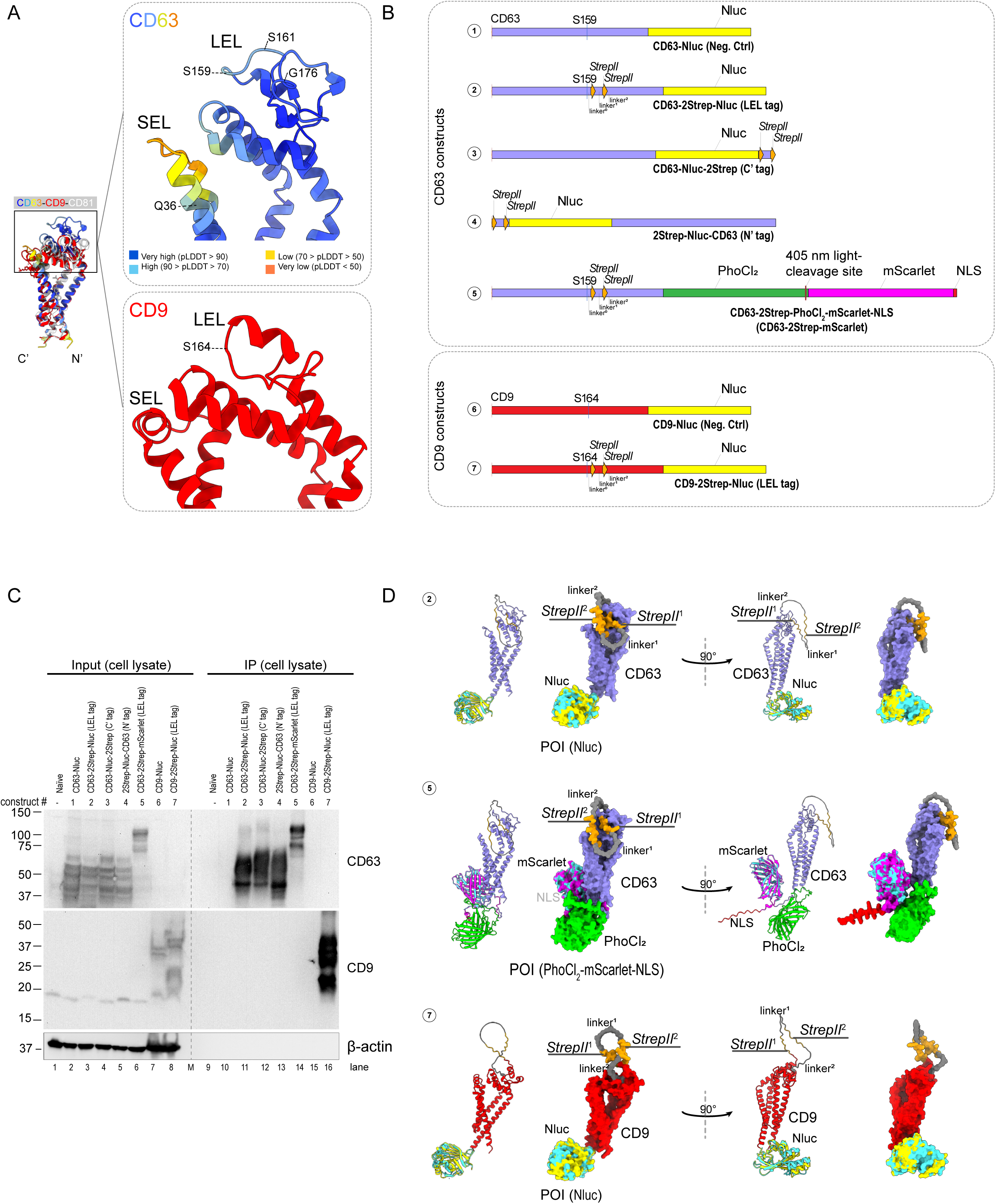
Construction and characterization of internally twin-Strep tagged CD63 and CD9 fused C-terminally with proteins of interest (POIs). (A) Twin-Strep tag insertion sites in the large extracellular loop (LEL) of CD63 and CD9. Left panel: Superimposed structures of high-confidence AlphaFold-predicted CD63 (AlphaFold Protein Structure Database: AF-P08962-F1-v4) with the crystal structures of CD9 (PDB: 6K4J, red) and cholesterol-bound CD81 (PDB: 5TCX, gray). Upper right: the three selected positions in the unstructured region of the LEL of CD63 (Serine 159, Serine 161, and Glycine 176) are denoted. The previously reported insertion site for GFP-pHluorin (Glutamine 36) in the small extracellular loop (SEL) is also marked. The predicted local distance difference test (pLDDT), which reflects the local confidence per residue in the AlphaFold prediction, is shown at the bottom. Lower right: the analogous CD9 insertion site in the unstructured region of the LEL after Serine 164 is shown. The superimposition was performed using the Matchmaker tool in UCSF ChimeraX [70]. (B) Schematics of CD63 and CD9 constructs bearing the twin-Strep (2Strep) tag at different positions. The twin-Strep consists of two tandem StrepII sequences; in CD63 construct #2 the 2Strep-tag is inserted into the LEL after Serine 159. The two StrepII motifs are each flanked by flexible linkers, as indicated. CD63-Nluc (construct #1), which lacks the 2Strep-tag, served as a negative control for affinity-based EV isolation. In construct #3 the 2Strep-tag is fused to the C-terminus of Nluc and in construct #4 to the N-terminus of CD63; both constructs were used as reference to evaluate how tag position affects CD63 protein integrity and capture efficiency. Construct #5 (CD63-2Strep-PhoCl_2_-mScarlet-NLS) includes a light-inducible PhoCl_2_ cleavage site and a C-terminal nuclear localization signal (NLS) on mScarlet. The analogous CD9 2Strep insertion in the LEL after Serine 164 (construct #7) and the corresponding CD9 control lacking the 2Strep-tag (construct #6) are also shown. Short names for each construct, used throughout the text, are given below each schematic. (C) Naïve HEK293T cells and cells stably expressing various CD63 and CD9 constructs were lysed using 1% Triton-X 100-containing buffer and analyzed by Western blot using CD63- and CD9-specific antibodies. Cell lysates were also subjected to Strep-TactinXT magnetic-bead isolation to capture 2Strep-tagged CD63 and CD9 and to examine the protein patterns resulting from different 2Strep-tag insertions in CD63. β-actin were used as loading controls. (D) Ribbon diagrams and space filling models of the proteins encoded by constructs (2), (5) and (7), as predicted by AlphaFold 3 [35]. In this prediction, the 2Strep-tag (orange) is exposed with no steric hindrance. Nluc and mScarlet are superimposed with their respective high-resolution crystal structures: Nluc at 1.95 Å (cyan; PDB entry: 5IBO) and mScarlet at 1.47 Å (cyan; PDB entry: 5LK4). Models are color-coded to match the construct schematics shown in Fig. 1B.

For further characterization, we selected the S159 insertion site for CD63, because this construct demonstrated high reproducibility and yielded approximately 40% of the input Nluc counts in the eluates (Fig. S1B). We further validated the protein expression of S159-inserted 2Strep-CD63 in cell lysates by western blot. As additional controls to evaluate the effect of 2Strep-tag insertion at this site in the LEL, we included CD63 constructs bearing the 2Strep-Nluc at the C–terminus (Fig 1B, construct #3) or N–terminus (construct #4). All 2Strep-tag CD63 variants showed comparable expression levels and overall banding patterns in cell lysates (Figure 1C, lanes 2–6). Moreover, after 2Strep-tag pulldown background was reduced (Figure 1C, lanes 11-14), indicating that 2Strep-tag insertion in the LEL permits proper CD63 expression and processing and allows efficient affinity-based capture.

In agreement with the experimental data, Alphafold3 prediction [35] of the S159-2Strep-tagged CD63 constructs (denoted as CD63-2Strep-Nluc and CD63-2Strep-mScarlet henceforth) suggested that the tag is exposed for EV capture and the CD63 domain well separated from the POI domains (Figure 1D). To further expand this engineering concept, we replicated our approach with CD9, another well-established tetraspanin enriched on small EVs [36, 37]. We inserted the 2Strep-tag into the unstructured region of the CD9 LEL after serine residue 164, analogous to the serine 159 insertion in CD63, and fused the protein C-terminally with the Nluc reporter (Fig. 1A, lower right; Fig. 1B, construct #7). This CD9-2Strep-Nluc construct was also well expressed (Fig. 1C, compare lanes 7 and 8) and efficiently captured from cell lysates (Fig. 1C, lanes 15-16).

### Subcellular localization of 2Strep-tagged CD63

CD63 is a crucial component of the endosomal pathway as it influences the sorting and packaging of molecules into multivesicular bodies (MVBs) for lysosomal degradation or exosome formation. Thus, CD63 has diverse functions in intracellular trafficking with the LEL being important for recruiting CD63 into EVs [38]. To validate proper subcellular distribution of CD63 tagged at position S159, we visualized the CD63-2Strep-mScarlet protein in the producer cells by using direct fluorescence imaging. We found that the mScarlet signals largely overlapped with Lamp1, a marker for endosomes and intraluminal vesicles that are EV precursors (Fig. 2A). This pattern was also observed for wild-type CD63 in naïve cells, indicating proper subcellular localization of the S159-tagged CD63.

**Figure 2.**
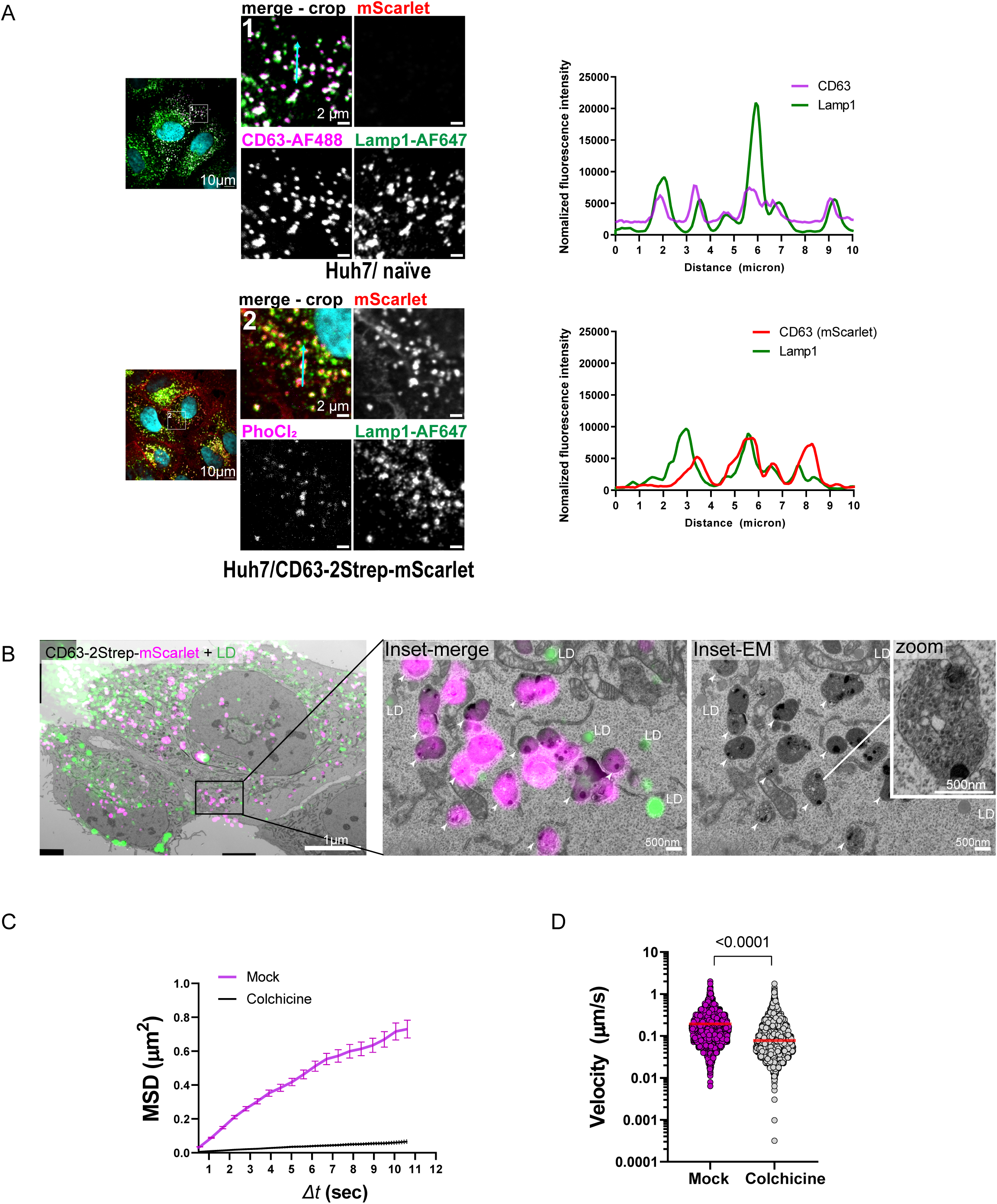
**Subcellular localization and trafficking of CD63-2Strep** (A) Colocalization of CD63-2Strep-mScarlet with the endosomal marker Lamp1. Naïve Huh7 cells and Huh7 cells stably expressing CD63-2Strep-mScarlet were fixed, immunostained for CD63 and Lamp1, and analyzed by confocal microscopy. Boxed areas in the left overview panels are shown as enlarged views in the panels on the right. Profiles on the right of the merged images were taken along the lines indicated by cyan arrows. The Alexa Fluor dyes (AFs) used to label the indicated proteins are shown. (B) Endosomal localization of CD63-2Strep-mScarlet. Huh7 cells stably expressing CD63-2Strep-mScarlet were analyzed by CLEM, using stained lipid droplets as fiducial markers to correlate electron and fluorescence micrographs. Left: merged overview of the CLEM micrograph. Middle: magnified inset of an area from the merged overview micrograph showing the correspondence of mScarlet signals to endosomes (arrowheads). Right: same magnified inset of the electron micrograph; endosomes are marked with arrowheads. One endosome is further magnified (top right) revealing numerous intraluminal vesicles contained within. Lipid droplets (LD) were stained with LipidTOX. (C) Microtubule-dependent trafficking of CD63-2Strep-mScarlet. Huh7 cells stably expressing CD63-2Strep-mScarlet were either mock-treated or treated with 80 µM colchicine for 1 hour to depolymerize microtubules. Cells were then analyzed by live-cell spinning-disk confocal microscopy, capturing images at 0.56-second intervals for a total duration of 1 minute. The dynamics of CD63-2Strep-mScarlet trafficking are represented by the mean squared displacement (MSD) of mScarlet signals. (D) Trafficking velocities of CD63-2Strep-mScarlet structures from (C). Statistical significance was determined using a two-tailed Mann-Whitney test.

To corroborate this assumption, we employed correlative light and electron microscopy (CLEM) by using lipid droplets as fiducial markers. We found that the mScarlet signals predominantly corresponded to endosomes (Fig. 2B, left and middle panels), which contained abundant small intraluminal vesicles (Fig. 2B, right panel, zoom). Since endosomes traffic in hepatocytes via the microtubule network [2, 39], we confirmed this trafficking pathway of CD63-2Strep-mScarlet using the microtubule-depolymerizing agent colchicine. Live-cell imaging analysis demonstrated microtubule-dependent movement of mScarlet signals, as colchicine-treated cells showed disrupted trafficking, evidenced by a reduction in both mean squared displacement (the average squared distance mScarlet signals traveled over time) and velocity (Fig. 2C and 2D; Video 1). Taken together, these data suggest that the insertion of the 2Strep-tag into the LEL of CD63 after serine residue 159, along with the flanking linkers, allows proper recruitment of CD63 into endosomes, ensures efficient secretion, and, most importantly, enables rapid and efficient isolation of CD63-containing EVs.

### Affinity-based isolation and characterization of 2Strep-tagged extracellular vesicles

To establish a rapid and minimally labor-intensive procedure for consistently obtaining highly pure EV preparations, we optimized the isolation protocol (Fig. 3A). Huh7 cells stably expressing CD63-2Strep-Nluc and control cells expressing non-tagged CD63-Nluc were cultured in EV-depleted media for 3 days to allow EV accumulation. Collected supernatants were filtered through 450-nm filter cups to remove dead cells and cell debris, and filtrates were concentrated using Centricon columns while simultaneously removing free proteins with a molecular weight below 100 kDa (Fig. 3B). We intentionally omitted the differential centrifugation steps (2,000, 4,000, and 10,000 *x g)* commonly used to remove large EVs, to assess the entire EV population that can be isolated with our Strep-based method. The concentrated supernatants were split and each part was used for EV isolation using two methods in parallel: the precipitation-based method employing a commercial total exosome isolation reagent, and the Strep-based method; the latter requiring no more than two hours, in contrast to the overnight incubation required for the precipitation-based method (Fig. 3B). As shown in Fig. 3C, the Strep-based approach achieved high enrichment of CD63-2Strep-Nluc with the Nluc signals being about three orders of magnitude above the background in the eluates and ∼60 times higher than the negative control. The recovery rate of EVs isolated by the Strep-based method was 72.4 ± 11.8% of the input (Fig. 3D).

**Figure 3.**
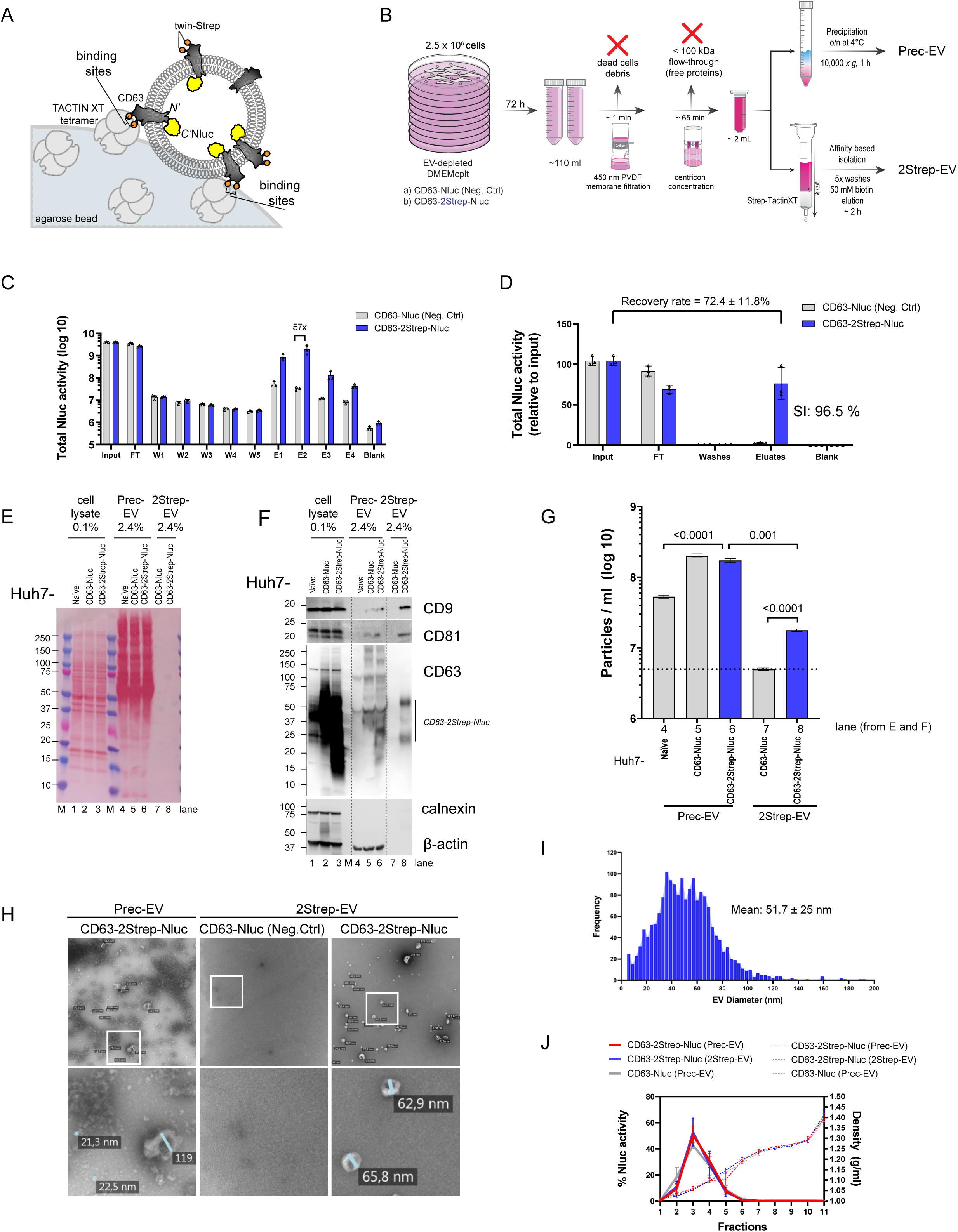
**Isolation of engineered extracellular vesicles using the Strep-based method** (A) Schematic depicting the binding of 2Strep-tagged CD63 on an EV (loaded with luminal Nluc) to the two binding pockets of the Strep-TactinXT tetramer immobilized on an agarose bead. (B) Schematic of the EV isolation procedure. Cell-conditioned media (∼110 mL) from Huh7 cells cultured in EV-depleted medium and stably expressing CD63-2Strep-Nluc or from cells expressing CD63-Nluc without a 2Strep-tag (negative control; Neg. Ctrl) were preprocessed by membrane filtration (cut-off 450 nm) to remove floating cells and debris. Proteins in the filtrates were removed by ultrafiltration (cut-off 100 kDa) and concentrated media were split for EV isolation using two methods: total EV isolation via precipitation (Prec-EV) using a commercial Total Exosome Isolation Reagent and twin-Strep tag-based affinity selection using the Strep-TactinXT approach (2Strep-EV). (C) Total Nluc activity contained in all fractions from the affinity-based EV isolation method, as described in (B). FT, flow-through; W, washes; E, eluates. Values were calculated for the total volume in each fraction and are displayed on a logarithmic scale. Data represent means (SD) from three biologically independent experiments. (D) Nluc activity in the various fractions from (C), shown relative to the input on a linear scale. The recovery rate and the specificity index (SI) are given. (E) Low protein complexity of affinity-captured 2Strep-EVs. Equivalent amounts of EVs isolated as described in (B), along with EVs from naïve Huh7 cells prepared by the precipitation-based method (Prec-EV), were analyzed by SDS-PAGE. After transfer to a PVDF membrane, proteins were stained with Ponceau S to visualize protein complexity prior to Western blot analysis. Cell lysates were loaded in parallel as references. M, protein molecular weight markers. Numbers refer to kDa. (F) Western blot analysis of the membrane from (E) using EV marker antibodies specified on the right. Calnexin, an ER-resident protein, served as an EV-negative marker; β-actin served as a marker commonly detected in total EVs isolated by the precipitation method. (G) Nanoparticle tracking analysis (NTA) of isolated EVs examined in (F). Data represent means (SEM) from three biologically independent experiments. Statistical significance was determined using a two-tailed Mann-Whitney test. (H) Transmission electron microscopy analysis after negative staining of EVs isolated by the precipitation-based method (Prec-EV) or by 2Strep tag-based affinity capture (2Strep-EV) as examined in (F). The boxed areas in the representative upper overview images are magnified in the corresponding lower panels. Lines indicate the diameters of representative vesicles. (I) Sizes of isolated 2Strep-EVs. Electron micrographs of isolated 2Strep-EVs in (H) were segmented using the Ilastik software and diameters were determined. A histogram showing vesicle diameters in nm is shown. Mean diameter (SD) = 51.7 ± 25 nm. (J) Density profiling of isolated EVs. Conditioned media of Huh7 cells expressing CD63 constructs specified on the top were used to isolate EVs by precipitation (Prec-EV) or via the 2Strep-tag (2Strep-EV) as described in (B). EV preparations were subjected to flotation ultracentrifugation using a 10-50% iodixanol gradient. Each fraction was analyzed for Nluc activity and values are displayed as the percentage of Nluc activity relative to the total Nluc across all fractions. Dotted lines refer to the density in the fractions (g/mL). Data represent means (SD) from three biologically independent experiments.

To evaluate the specificity of 2Strep-tag based isolation method, we defined a specificity index (SI). The SI represents the ratio of Nluc activity in the eluate of the CD63-2Strep-Nluc sample, divided by the combined Nluc values from both CD63-2Strep-Nluc and the negative control lacking the 2Strep-tag (CD63-Nluc) in the eluate. A maximum value of 100% indicates complete tag-specific capture; in this setup, the SI was 96.5% (Fig. 3D).

We further examined the protein content of the EV fractions that were isolated by using SDS-PAGE, Ponceau staining of membrane-transferred proteins and Western blot analysis. While EV samples isolated by the chemical precipitation method (referred to as Prec-EVs henceforth) contained high amounts of proteins, likely due to FCS-derived and cell-secreted contaminants (Fig. 3E, lanes 4 - 6), the EVs isolated with the Strep-based method (2Strep-EVs) contained significantly less protein (Fig. 3E, lanes 7, 8). Importantly, Western blot analysis revealed bona fide EV markers in the 2Strep-EV preparation, including CD63, CD9, and CD81 (Fig. 3F, lane 8), which were absent in the untagged negative control sample (lane 7), indicating specificity of the isolation method. In addition, Prec-EVs displayed stronger smearing patterns of CD63, reflecting the isolation of complex EV populations that include both endogenous, untagged and 2Strep-tagged CD63, as well as aggregated CD63 (Fig. 3F, lanes 5, 6). While the EV preparations isolated with either approach tested negative for calnexin, an ER-resident protein whose absence is commonly used as purity control for EVs, only the EV preparation generated with the Strep-based method was devoid of beta-actin, which is frequently found in EV preparations isolated via precipitation [3] (Fig. 3F, lanes 4 - 8).

Nanoparticle tracking analysis (NTA) revealed a consistently lower particle count in the 2Strep-EVs compared to Prec-EVs, confirming reduced amounts of contaminant particles in the affinity-isolated EV preparation (Fig. 3G and S1C). Moreover, negative staining and transmission electron microscopy (TEM) analysis of 2Strep-EVs revealed abundant small EVs with an average diameter of ∼52 nm and a clean background, whereas EV preparations generated by the precipitation-based method contained numerous protein aggregates and lipoprotein-like structures (Fig. 3H, 3I).

To further characterize 2Strep-tag isolated EVs, we employed flotation gradient centrifugation to examine their density profiles. EVs containing CD63-2Strep-Nluc isolated by precipitation or the 2Strep-based method, and EVs precipitated from culture supernatants of cells expressing non-tagged CD63-Nluc, exhibited similar density profiles in the range of ∼1.05-1.1 g/mL (Fig. 3J), arguing that 2Strep-EVs remain structurally intact. We further tested the feasibility of the isolation approach using cell-conditioned media devoid of FCS. Purity of the 2Strep-EV preparation could be increased further by culturing the cells in FCS-free media for 24 hours prior to supernatant harvest. Isolated 2Strep-EVs produced under this condition reached a SI index of 99.0% (Fig. S1D), which is in good agreement with the TEM analysis (Fig. S1E). Taken together, these results show that the 2Strep-based method enables reproducible and highly selective enrichment of engineered EVs while ensuring efficient removal of contaminants.

### Proteome analysis of affinity-isolated 2Strep-tagged extracellular vesicles

To investigate the proteome profile of 2Strep-tag-isolated EVs, we performed mass spectrometry analysis of the EVs characterized in Fig. 3. As expected, 2Strep-purified EVs contained significantly lower number of proteins than Prec-EVs (Fig. 4A). Importantly, despite this reduced overall protein content, the analysis revealed a largely comparable enrichment of conventional EV protein markers in 2Strep-EVs compared to Prec-EV preparations, including CD63, CD81, and CD9 (Fig. 4B). In addition, other common EV markers such as lactadherin, integrin β-1, ANXA2 as well as TSPAN3 and TSPAN6 were abundantly detected in both Prec-EV and 2Strep-EV samples. Interestingly, in 2Strep-EVs we detected additional putative EV-associated proteins, including other CD molecules such as tetraspanins, syntenins, flotillins, integrins, and annexins. Importantly, contaminants were markedly reduced in 2Strep-EVs compared to Prec-EVs, with a ≥97% depletion of FCS-derived and cell-secreted impurities. Representative impurities that were removed included nanoparticle-associated contaminants such as various apolipoproteins, FCS-derived EV proteins like CD molecules, heat shock proteins, integrins, heparan sulfate proteoglycan, and cell adhesion molecules (Fig. 4C). Other major contaminants being removed by the Strep-based method included bovine and human serum albumin, calnexin, and complement factors (Fig. 4D), in line with the Western blot analysis (Fig. 3F). We also identified EV-associated proteins not previously reported including lipolysis-stimulated lipoprotein receptor (LSR) as well as recently discovered proteins such as Prominin-1 (PROM1), Junction Plakoglobin (JUP), and Stomatin (STOM), among others (Fig. 4E). Taken together, CD63-2Strep-tagged EVs isolated by the Strep-based method were highly enriched for the engineered EVs. The profound depletion of impurities allowed the detection of low-abundance putative EV markers.

**Figure 4.**
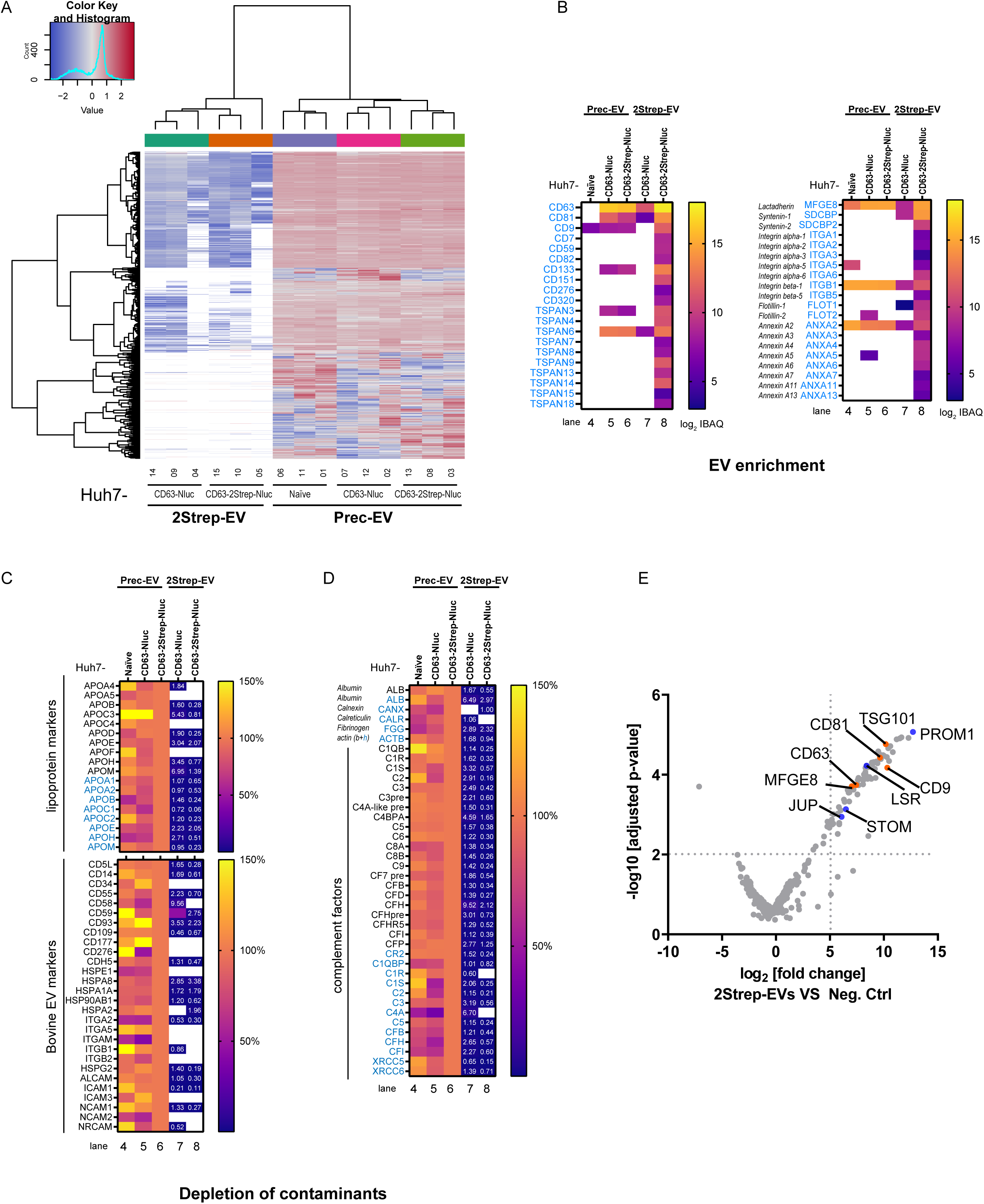
**Comparison of the proteome of extracellular vesicles isolated by the 2Strep- or the precipitation-based method** (A) Heatmap showing the Z-scores of proteins detected in EV preparations obtained by precipitation (Prec-EV) or 2Strep affinity-based capture (2Strep-EV). Prec-EV and 2Strep-EV samples, as described in Figure 3, were analyzed by LC-MS/MS and grouped using data-independent acquisition (DIA)-based clustering. Data were acquired from three independent biological replicates, with the sample order specified at the bottom (samples 1-15). (B) Selective enrichment of EV markers in 2Strep-isolated EVs. [Left panel] Selected EV marker proteins of human origin include the classical markers CD63, CD81, and CD9, along with other detected CD proteins and tetraspanins (TSPAN), identified by matching detected peptide sequences against the human reference proteome from UniProt. [Right panel] Additional representative EV-associated proteins, most of them only detectable in 2Strep-isolated EV samples because of higher purity. The mean IBAQ values of 3 independent biological replicates were log_2_-transformed and are shown on the heatmap. The lane numbers correspond to the samples in Fig. 3E and 3F. White blocks indicate no detection. (C) Depletion of contaminating nanoparticles by 2Strep affinity-based isolation. Selected lipoprotein markers (apolipoproteins, APOs) and bovine EV markers detected in the EV samples specified on the top are shown. The mean IBAQ values from 3 biologically independent replicates were log_2_-transformed. Amounts of each protein detected in EV preparations that were isolated by the precipitation-based method from Huh7 cells expressing CD63-2Strep-Nluc were set to 100%. Protein amounts are indicated by color coding and, in cases of low abundance (<10%), also by numbers. White blocks indicate no protein detection. Proteins of human origin are highlighted in blue. The lane numbers correspond to samples in Fig. 3E and 3F. (D) Depletion of major representative non-vesicular contaminants, including albumin, calnexin, calreticulin, fibrinogen, actin, and complement factors. (E) Volcano plot comparing 2Strep-EVs isolated from Huh7 cells expressing CD63-2Strep-Nluc using the Strep-based method (lane #8, Fig. 3E and 3F) with the negative control lacking the twin-Strep tag (lane #7). Protein hits with high log_2_ fold change and significant −log10(p) values, including representative EV marker proteins CD63, CD81, CD9, TSG101, and MFGE8 (orange dots) are identified. Potential novel EV proteins such as LSR (lipolysis-stimulated lipoprotein receptor), Prominin-1 (PROM1), Stomatin (STOM), and Junction Plakoglobin (JUP), are also highlighted (blue dots). Dotted lines indicate thresholds of log_2_FC ≥ 5 (fold change ≥ 32) and - log_10_(p) ≥ 2 (p ≤ 0.01).

### Efficient separation of 2Strep-tagged extracellular vesicles from virus particles

Engineered EVs for targeted delivery must be devoid of pathogenic viruses. To evaluate the performance of our isolation method, we tested its ability to separate CD63-2Strep-tagged EVs from virus particles (virions). As a model, we used the hepatitis C virus (HCV), because it replicates and spreads efficiently in Huh7 cells. Moreover, HCV virions most likely assemble at the ER and are transported via MVBs out of the cell, thus passaging through the same compartment as EVs [40–42]. Notably, HCV particles have similar density and size to EVs (density of ∼1.09 g/ml and mean size of ∼62 nm) [43, 44], making conventional separation of virions and EVs challenging. Huh7 cells stably expressing CD63-2Strep-Nluc, or the CD63-Nluc control construct, were infected with HCV for 3 days to allow productive viral replication and the release of virus progeny into the culture supernatant (Fig. 5A). Viral replication in these cells was confirmed by the detection of the viral proteins NS5A and capsid (core), the latter being part of virions (Fig. 5B). We isolated CD63-2Strep-Nluc containing EVs from culture supernatants of HCV infected cells by using the Strep-based method. High Nluc activity in the eluate confirmed the enrichment of CD63-2Strep-Nluc containing EVs (Fig. 5C) whereas virus particles in the same input were depleted by >99.5% (Fig. 5D). We conclude that our approach allows the separation of CD63-2Strep-tagged EVs from virus particles present in the same input material, thus guaranteeing high purity of the EV preparation and rigorous removal of viral contaminants.

**Figure 5.**
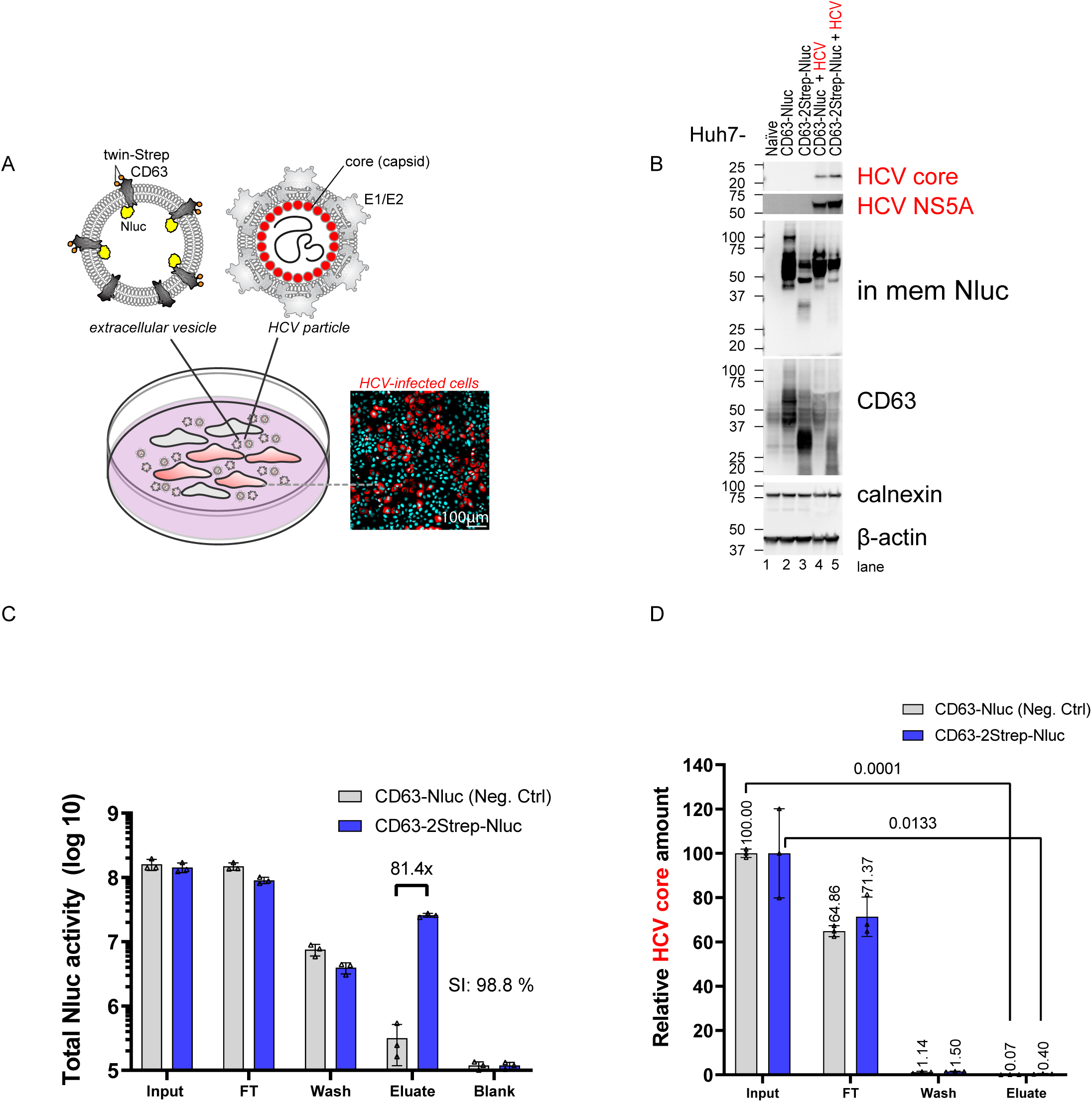
**Efficient separation of virus particles from EVs enriched by the Strep-based approach** (A) Schematic depicting Huh7 cells producing 2Strep-EVs and being in addition infected with HCV (red cells). Supernatants from these cells contain both EVs with CD63-2Strep-Nluc and HCV particles, the latter being decorated with the envelope glycoproteins E1 and E2 and containing a nucleocapsid composed of the core protein and the RNA genome (black line). In the light microscopy image, intracellular HCV non-structural protein 5A (NS5A), a marker of HCV replication, was detected by immunofluorescence (red) while nuclear DNA was counterstained with DAPI (cyan). (B) Huh7 cells stably expressing CD63-2Strep-Nluc and control cells expressing non-tagged CD63-Nluc (CD63-Nluc) were infected with HCV. At 72 hours post-infection, cell lysates were analyzed by Western blot using antibodies specified on the right. Lysates of uninfected control cells were loaded in parallel. CD63-Nluc activity was visualized using the in-membrane (in mem) Nluc activity assay. Calnexin and β-actin served as loading controls. (C) Efficient removal of HCV and enrichment of 2Strep-EVs from culture supernatants containing a mixture of virions and EVs. 2Strep-EVs present in HCV-containing media were isolated using the Strep-based method. Nluc activity was measured in all fractions of the isolation process, including the flow-through (FT), washes, and eluates. Values were calculated for the total volume in each fraction and are displayed on a logarithmic scale. Data represent means (SD) from three biologically independent experiments. SI: specificity index. (D) Quantification of HCV core antigen present in isolated 2Strep-EVs. HCV core protein levels in all fractions from (C) were quantified using the HCV Core CMIA assay. Values were calculated for the total volume of each fraction and normalized to the input that was set to 100. Data represent means (SD) from three biologically independent experiments. Statistical significance was determined using unpaired *t*-test with Welch’s correction.

### Cellular uptake of 2Strep-tagged extracellular vesicles

Considering the possible use of EVs as vehicle to deliver a given POI, cellular uptake of the EVs is an important prerequisite. Therefore, we assessed cellular uptake of 2Strep-EVs. In the first set of experiments, we used similar Nluc-input equivalents of 2Strep-EVs and Prec-EVs isolated from culture supernatant of Huh7-derived cells (Fig. 3). These EVs were added to naïve Huh7 cells and evaluated for cellular uptake at 2 and 6 hours post-incubation at 37°C (Fig. S2A and S2B). Incubation for 2 hours at 4°C was used to assess the background signal corresponding to EVs merely adsorbed to the cells. Uptake of Nluc contained in either EV preparation was comparable at 2 and 6 hours at 37°C (Fig. S2B).

Next, we aimed to visualize the uptake of 2Strep-EVs. However, the labeling of EVs with dyes often presents challenges, including incomplete labeling of EVs, non-specific labeling of non-EV particles, formation of dye micelles or dye-protein aggregates, and interference from autofluorescence of unlabeled vesicles [20, 21]. Therefore, we employed 2Strep-EVs containing the bright monomeric fluorescent protein mScarlet (Fig. 1B). We isolated these EVs from Huh7 cells using the 2Strep-based method (Fig. 3) and Western blot analysis confirmed the enrichment of mScarlet-containing EVs (Fig. 6A). To facilitate uptake analysis, we employed Huh7-derived recipient cells stably expressing eYFP that was fused with the farnesylation signal CaaX from the human HRAS protein to anchor eYFP to the plasma membrane and to endosomes. We incubated these recipient cells with isolated mScarlet-containing 2Strep-EVs or control Prec-EVs from naïve Huh7 cells for 6 hours and monitored the internalized mScarlet signals within the cell boundaries by live-cell confocal imaging to avoid fixation artifacts (Fig. 6B). We detected mScarlet signals within the cell boundaries, partially colocalizing with eYFP-labeled vesicular structures, most likely corresponding to endosomes, which are the primary sites of internalized EVs (Fig. 6C and 6D, Video 2).

**Figure 6.**
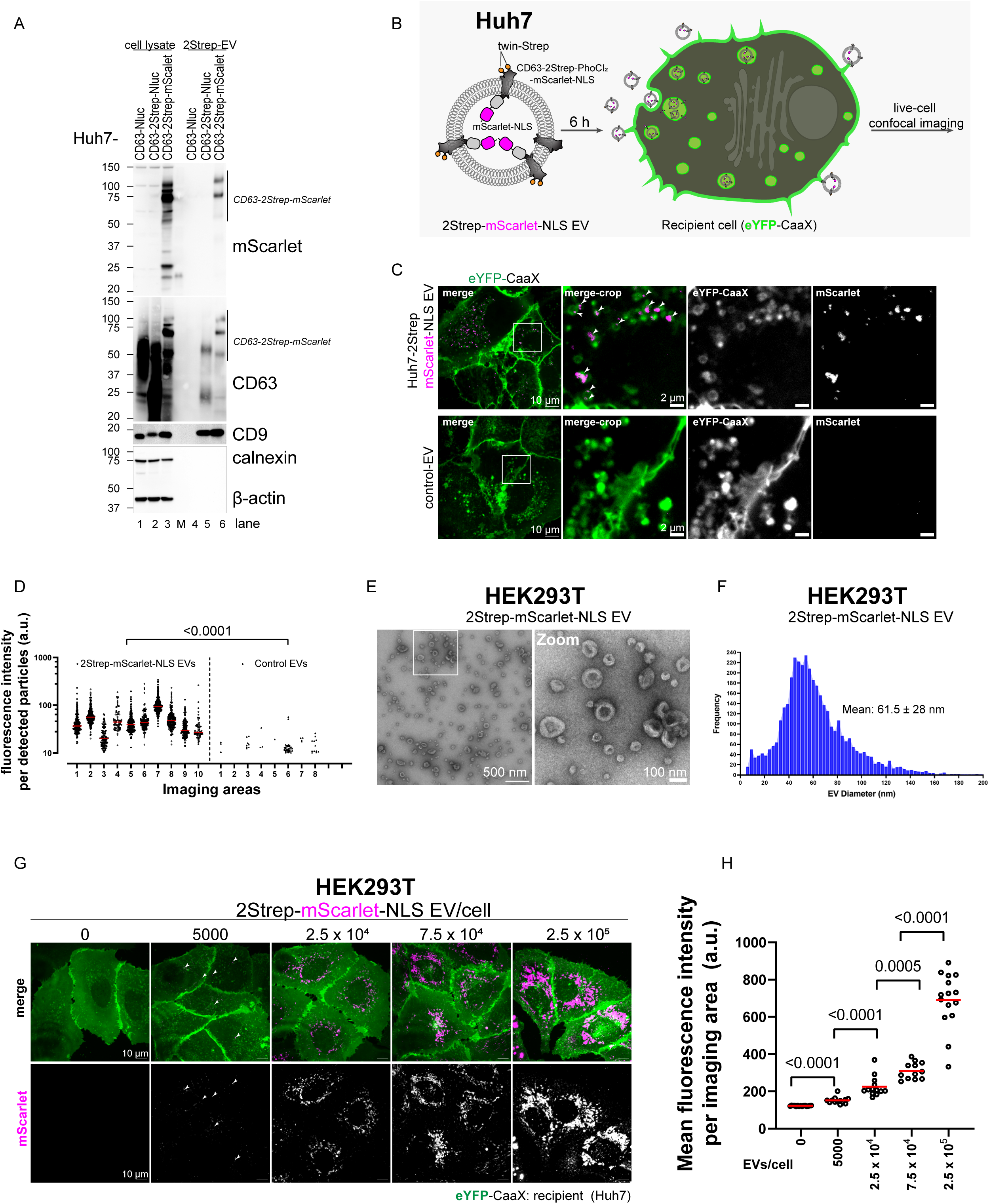
**Uptake of 2Strep-tagged extracellular vesicles encapsulating a protein of interest** (A) Western blot analysis of Huh7 cells expressing CD63-2Strep-PhoCl_2_-mScarlet-NLS (referred to as CD63-2Strep-mScarlet for simplicity) and of EVs isolated from the supernatant of these cells by the 2Strep-based method. Samples were analyzed by Western blot using an RFP-specific antibody or EV marker antibodies specified on the right. As a control, 2Strep-EVs containing CD63-Nluc (refer to Fig. 3) were included. CD63-Nluc without 2Strep-tag served as negative control. Calnexin and β-actin served as loading controls. (B) Schematic of the experimental procedure used for EV uptake analysis. Equivalent amounts of affinity-isolated 2Strep-EVs encapsulating mScarlet-NLS or control EVs isolated by precipitation (Prec-EVs) from the culture supernatant of naïve Huh7 cells, were added to recipient cells. Recipient cells express an eYFP-CaaX fusion protein (CaaX refers to the farnesylation sequence derived from the human HRAS protein) to visualize cellular membranes, including the plasma membrane and endosomes [2]. At 6 hours post-incubation, cells were subjected to live-cell confocal imaging to detect internalized fluorescent signals. (C) Internalization of 2Strep-EVs encapsulating mScarlet-NLS. Fluorescent signals from internalized mScarlet associated with vesicular structures are shown. The boxed areas in the overview images (left) are magnified in the right panels, highlighting internalized mScarlet puncta (arrowheads). (D) Quantification of internalized fluorescent foci from (C). Each dot represents one detected particle from one imaging field of view (125 x 161 microns), with the relative signal intensity above the threshold shown on the Y-axis. Statistical significance was determined using a two-tailed nested *t*-test. (E) Isolated 2Strep-EVs encapsulating mScarlet-NLS, derived from HEK293T cells expressing CD63-2Strep-mScarlet, were visualized by negative staining and TEM. The boxed area in the overview electron micrograph (left) is magnified in the right panel, showing highly enriched EVs of small sizes. (F) Sizes of isolated 2Strep-EVs encapsulating mScarlet-NLS. Isolated EVs in electron micrographs from (E) were segmented using the Ilastik software and diameters were determined. A histogram showing vesicle diameters in nm is shown. Mean diameter (SD) = 61.5 ± 28 nm. (G) Dose-dependent uptake of 2Strep-EVs encapsulating mScarlet-NLS, derived from HEK293T cells. Huh7-derived cells expressing the membrane marker eYFP-CaaX were incubated with increasing EV amounts specified on the top of the panels for 24 hours. Internalized EVs were analyzed by live-cell imaging covering a 2 µm Z-stack. Maximum Z-projection images show internalized EV signals within the boundaries of recipient cells. Arrowheads indicate representative foci of internalized mScarlet signals that appear as discrete puncta in the low-dose inoculated cells. (H) Quantification of internalized mScarlet signals from (G). Each dot represents the mean fluorescence intensity of a maximum Z-projected imaging view (125 x 161 microns). Statistical significance was determined using a two-tailed Mann-Whitney test.

The EV preparations described so far have been obtained from human hepatoma cells (Huh7). Since HEK293T cells are widely used for EV production, we evaluated the versatility of our EV production and purification approach by isolating 2Strep-EVs from conditioned media of HEK293T cells stably expressing CD63-2Strep-Nluc. EVs produced in these cells could be enriched by affinity isolation as rapidly and efficiently (specificity index of > 99%) as EVs derived from Huh7 cells (Fig. S2C). Likewise, CD9-2Strep EVs could be captured from culture supernatant of HEK293T cells with an efficiency comparable to CD63-containing engineered EVs (Fig. S2D), demonstrating the versatility of this tagging approach. This result aligns with the predicted structure of CD9-2Strep-Nluc, where the 2Strep-tag is expected to be exposed (Fig. 1D).

Also mScarlet-containing EVs, released from HEK293T cells stably expressing CD63-2Strep-mScarlet, could be enriched with high yield and purity (Fig. 6E and S2E). Purified EVs were small with a mean diameter of 61.5 ± 28 nm (Fig. 6F). Incubation of Huh7 recipient cells stably expressing eYFP-CaaX with these EVs at varying doses demonstrated specific uptake by recipient cells. Uptake signals became detectable at an NTA-based dose of 5,000 EVs/cell (corresponding to 5 fg CD63/cell) by both Western blot analysis (Fig. S2F) and live-cell imaging (Fig. 6G and 6H). These results suggest that 2Strep-EVs isolated by our Strep-based method are competent for endocytic uptake.

### Cytoplasmic release of proteins of interest contained in engineered 2Strep-isolated extracellular vesicles

We next tested whether the mScarlet reporter protein contained in 2Strep-isolated EVs could be released after uptake. Since natural EVs do not carry fusogenic proteins, we aimed to enhance EV fusion with endosomal membranes to facilitate cargo release. To achieve this, 2Strep-tagged EVs were coated with the vesicular stomatitis virus glycoprotein (VSV-G) [23, 45–47]. HEK293T cells stably expressing CD63-2Strep-mScarlet were transiently transfected with a VSV-G construct and EVs were isolated using the Strep-based method (Fig. 3). Isolated EVs were illuminated at 405 nm to induce PhoCl_2_-mediated release of mScarlet into the EV lumen prior to addition to recipient cells (Fig. 7A).

**Figure 7.**
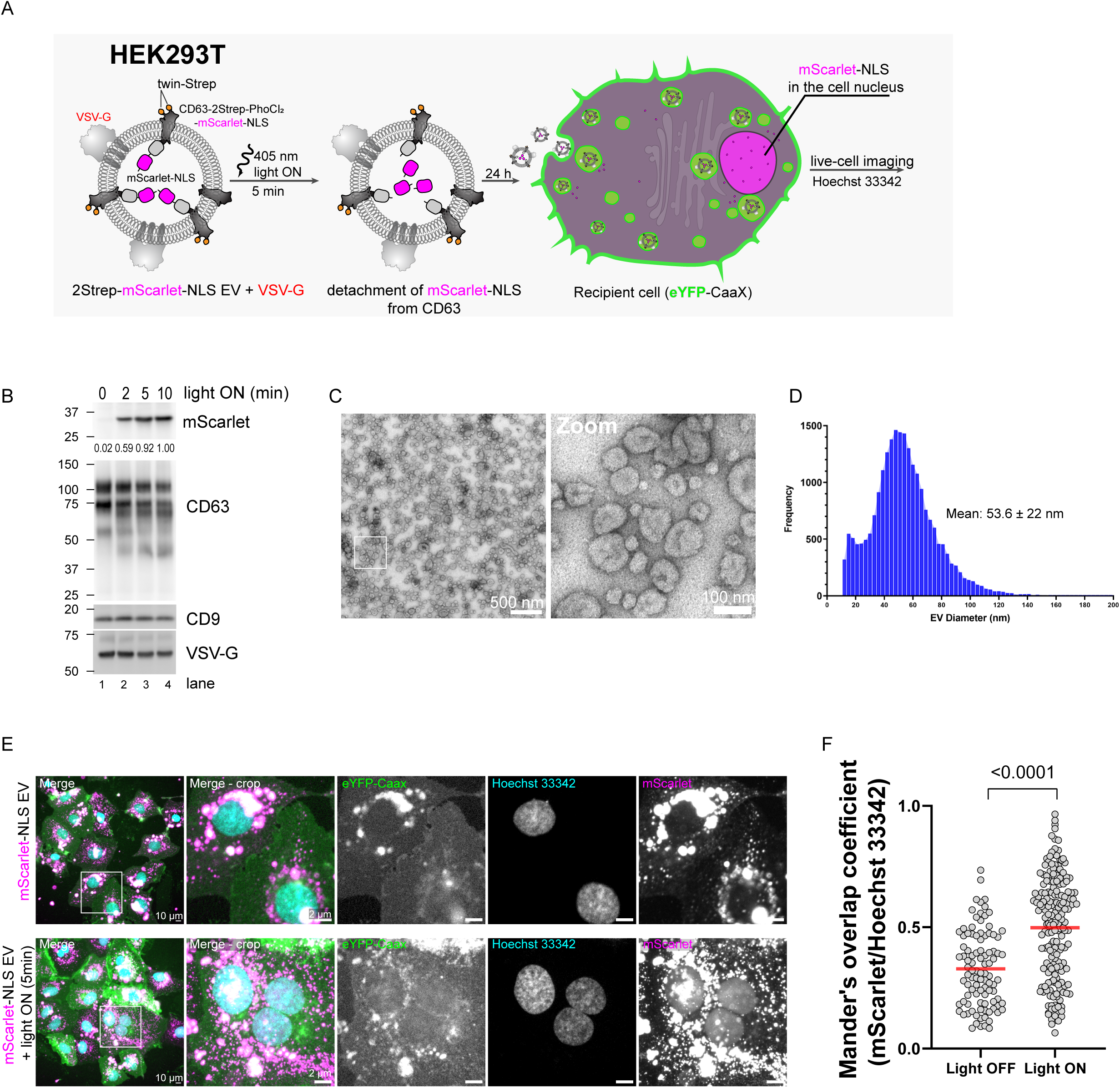
**Endosomal escape of delivered proteins following cellular uptake of isolated twin-Strep-tagged extracellular vesicles** (A) Schematic of the experimental procedure used for mScarlet-NLS endosomal escape analysis. HEK293T cells expressing CD63-2Strep-PhoCl_2_-mScarlet-NLS (referred to as CD63-2Strep-mScarlet for simplicity) were transiently transfected with a VSV-G expression plasmid to enhance EV fusogenicity. Affinity-isolated 2Strep-EVs from the culture supernatant of these cells were either left untreated or illuminated with 405 nm light to induce PhoCl_2_ cleavage, thereby releasing mScarlet-NLS from CD63 within the EV lumen. These EVs were then added to recipient cells expressing the eYFP-CaaX membrane marker. After 24 hours of incubation, cells were counterstained with Hoechst 33342 to visualize cell nuclei and subjected to live-cell imaging to detect internalized mScarlet-NLS signals and assess their nuclear localization. Endosomal release was determined by quantifying mScarlet-NLS signals in cell nuclei. (B) Western blot analysis of affinity-isolated VSV-G-coated 2Strep-EVs encapsulating mScarlet-NLS, derived from HEK293T cells. EVs were either left untreated or treated with 405 nm light for 2, 5, or 10 minutes. Samples were analyzed by Western blot using an RFP-specific antibody to detect mScarlet or a CD63-specific antibody. CD9 and VSV-G served as cleavage and loading controls. (C) Affinity-isolated VSV-G-coated 2Strep-mScarlet-NLS EVs were visualized by negative staining and TEM. The boxed area in the overview electron micrograph (left) is magnified in the right panel, showing highly enriched EVs of small sizes (<200 nm). (D) Sizes of VSV-G-coated 2Strep-mScarlet-NLS EVs. Electron micrographs of isolated EVs in (C) were segmented using the Ilastik software, and diameters were determined. A histogram showing vesicle diameters in nm is shown. Mean diameter (SD) = 53.6 ± 22 nm. (E) Endosomal escape of mScarlet-NLS following internalization of VSV-G-coated 2Strep-mScarlet-NLS EVs. Recipient cells were incubated with EVs at a dose of 1.6 x 10^4^ EVs/cell (as described in Fig. 7A). Boxed regions in the overview images (left) are shown at higher magnification in the right panels. Magnifications highlight internalized mScarlet-NLS signals (upper panel) and additional diffuse nuclear mScarlet signals (lower panel) in cells that received EVs illuminated to release mScarlet-NLS from the CD63 scaffold prior to addition. (F) Colocalization analysis of mScarlet signals with nuclear staining. Nuclei visualized by Hoechst staining were used for primary object detection, and the Mander’s overlap coefficient of mScarlet signals over Hoechst signals within the identified objects was calculated using the CellProfiler software [71]. Data are presented as mean (SD), with individual dots representing single cells.

Western blot analysis confirmed enrichment of 2Strep-mScarlet EVs along with VSV-G and light-induced PhoCl_2_ cleavage releasing the ∼26.5 kDa mScarlet protein with the C-terminal nuclear localization signal (NLS) (Fig. 7B). Negative staining and TEM analysis further validated the morphology of the isolated EVs, with a mean diameter of 53.6 ± 22 nm (Fig. 7C, D). After 24 hours of incubation of the cells with these EVs, live-cell imaging revealed well-detectable uptake, indicated by multiple mScarlet foci within the boundaries of recipient cells (Fig. 7E, upper panel). Importantly, when EVs were pretreated with light, mScarlet-NLS was deposited in cell nuclei, confirming successful endosomal escape (Fig. 7E, lower panel; Fig. 7F).

## Discussion

Achieving selective isolation of intact engineered EVs with sufficiently high purity and yield is a fundamental prerequisite for the study of EV biology and EV-based translational applications. The majority of published studies still rely on bulk EV preparations that are of limited selectivity for engineered versus native EVs and frequently contaminated with non-vesicular nanoparticles, cell-and serum-derived components, or virus-like particles. Such impurities confound engineered EV dosing, obscure mechanistic interpretation, and raise concerns regarding functional assignment, as observed biological effects may originate from co-isolated contaminants rather than from engineered EVs themselves. Addressing this fundamental limitation, we sought to introduce one of the most advanced affinity purification technologies into the EV field, thereby providing the research community with a robust means to isolate engineered EVs with exceptional selectivity, integrity, and reproducibility under native conditions. The central conceptual advance of our work is the structure-guided internal placement of the 2Strep-tag within canonical EV-associated tetraspanins. As shown for CD63, the tagged protein retains biological activity as it supports EV biogenesis and retains EV integrity as well as cargo functionality.

CD63 is one of the most widely used EV scaffolds owing to its robust incorporation into EVs and its strong capacity for luminal cargo loading [28]. However, tetraspanin topology, with both N- and C-termini oriented toward the EV lumen (Fig. 3A), has fundamentally limited the applicability of conventional 2Strep affinity tagging strategies. As a result, high-affinity peptide-ligand systems such as Strep technology, despite their widespread use for native protein complex purification, have remained inaccessible for EV isolation. Here, we overcome this standing constraint by inserting the 2Strep-tag into an unstructured region of the CD63 large extracellular loop (LEL), flanked by flexible linkers. This internal tagging strategy exposes the affinity tag on the EV surface without perturbing CD63 trafficking, localization, or EV sorting, thereby converting CD63 into a dual-functional scaffold that simultaneously enables surface-based affinity capture and luminal cargo loading. This design unlocks the use of Strep-TactinXT, a directed-evolution–derived streptavidin variant with near-covalent affinity and exceptional specificity for the 2Strep-tag [24–26]. The system offers several decisive advantages: (i) antibody-free capture, eliminating antibody contamination; (ii) exceptionally low nonspecific binding, allowing extensive washing without EV loss; and (iii) mild, fully reversible elution via biotin competition under physiological buffer conditions. Together, these properties enable non-destructive isolation of intact EVs at an excellent level of specificity and purity.

Our strategy enables the selective isolation of engineered EVs containing protein cargo of interest, achieving yields of ∼40–80% and ∼97% purity from complex mixtures containing non-engineered EVs, cell-secreted impurities, serum-derived components (Fig. S1F), and even virus particles with EV-resembling biophysical properties. The entire capture and elution process is completed within two hours using a gravity-flow column setup, eliminating the need for complex equipment such as ultracentrifuges or advanced chromatography systems, thereby making the method applicable to a wide range of laboratories. For future upscaling of EV production, including GMP-compliant workflows aimed at clinical applications and reduced plastic use, appropriate suspension cell lines, such as GMP-banked HEK293-F with detailed lineage history, and grown in serum-free, chemically defined media, will be beneficial. A Strep-TactinXT FPLC column setup would be well-suited for integration into automated, scalable purification pipelines. While our current strategy is highly effective, further optimization might be required for such upscaling and could be integrated with other isolation approaches to further boost the purity of the final EV preparation. Nevertheless, the method reported here should provide an important starting point towards this direction.

The high purity of affinity-isolated EVs enabled sensitive analysis by mass spectrometry. Apart from the canonical EV markers (CD63, CD81, CD9), we detected low-abundance proteins in affinity-isolated EVs without mass spec system saturation. These proteins included the lipolysis-stimulated lipoprotein receptor (LSR), a protein complex in the liver that provides binding sites for ApoE and ApoB upon free fatty acid engagement [48, 49]. Other examples of uncommon EV-associated proteins are Prominin-1 (PROM1) [50], Stomatin (STOM) [51] and Plakoglobin (JUP) [52], which are potential candidates for novel EV markers. However, confirmation and functional characterization of these EV-associated proteins remain topics for future studies.

One major barrier to the therapeutic application of EVs is the lack of standardized dosing [3, 16]. Inaccurate dosing often arises from quantifying EV numbers in impure preparations, which does not reflect the real dose of functionalized EVs. Because our method selectively enriched engineered EVs, we could determine particle numbers and vesicle size without interference from abundant nanoparticles derived from cells, serum, or protein aggregates. We observed that CD63-positive EVs produced in Huh7 and HEK293T cells included a substantial population of small vesicles below 80 nm diameter, in addition to the expected 80-150 nm diameter range. These smaller vesicles fall below the optimal detection threshold of NTA [53], suggesting that dosing based on NTA particle counts largely underestimates the true number of EVs. Nevertheless, quantifying highly pure EVs also has challenges [3]. Notably, the total protein content of 2Strep-tagged EVs isolated from Huh7 cells was often below the linear detection range of commonly used protein assays such as Bradford or BCA. In this regard, using a highly sensitive nano-flow cytometer capable of detecting EVs down to 40 nm could increase the detection range for small EVs. Alternatively, here we combined ELISA with NTA to quantify the numbers of EVs isolated from HEK293T cells, but we acknowledge that a more accurate approach to determine EV numbers remains elusive.

To validate the functionality of engineered CD63 and CD9 in Huh7 and HEK293T cells, we selected Nluc and mScarlet (the latter only for CD63) as POIs. These reporters enabled monitoring and quantification of the isolation process as well as tracking of cellular uptake of engineered EVs, and they can be potentially applied to many cell systems of interest. The delivery of endogenously loaded functional proteins within the EV lumen has been successfully demonstrated in previous studies using both cell culture and murine models. Notable examples of delivered proteins include Cas9 for genome editing, catalase for mitigating oxidative stress, and transcription factors such as NF-κB for suppressing systemic inflammation [23, 33, 34, 47]. Our approach not only preserves POI integrity via encapsulation within the EV lumen, but also allows optogenetically controlled POI release, combined with the selective isolation of engineered EVs via surface-expressed 2Strep-tagged CD63, which ensures high enrichment of the desired EVs.

## Conclusions

We report a structure-guided affinity tagging strategy that enables rapid, highly selective, antibody-free, and non-destructive isolation of intact engineered EVs. By internally inserting a twin-Strep tag into unstructured regions of the extracellular loops of the canonical tetraspanins CD63 and CD9, we overcome a fundamental topological limitation that has previously precluded the application of the Strep technology to intact EVs. This design converts tetraspanins into dual-functional scaffolds that support surface-accessible affinity capture under native conditions and at the same time, by C-terminal fusion with a protein-of-interest, efficient luminal cargo loading.

The resulting affinity-based workflow yields engineered EVs with exceptional specificity and recovery, enabling rigorous separation from non-engineered vesicles, cell- and serum-derived nanoparticles as well as virus-like particles. The high integrity of the isolated EVs enables functional delivery of EV luminal cargo, including optogenetically controlled cargo release, and enables sensitive proteomic analyses of engineered EVs without interference from abundant contaminants. Owing to its modular design, this platform is readily extendable to other EV-associated scaffolds and thus provides a potentially universal strategy for the efficient isolation of engineered EVs encapsulating functional POIs.

## Materials and Methods

### Materials

#### Antibodies and reagents

The following antibodies and reagents were used in this study: rabbit CD63 antibody (Cell Signaling, 52090S); rabbit RFP antibody (Rockland, 50961); mouse CD63 antibody-Alexa Fluor 488 (Santa Cruz, sc-5275); mouse Strep-tag II antibody (IBA Lifesciences, 2-1507-001); rabbit CD9 antibody (Cell Signaling, 13174S); mouse HCV NS5A 9E10 antibody (kind gift from Charles Rice, Rockefeller University); rabbit HCV core C830 antibody [54]; rabbit Calnexin C5C9 antibody (Cell Signaling, 2679S); mouse β-actin antibody (Sigma Aldrich, A5441); Strep-TactinXT DY-549 (IBA Lifesciences, 2-1565-050); MagStrep Strep-TactinXT beads (IBA Lifesciences, 2-5090-010).

#### Materials for affinity-based extracellular vesicles isolation

For supernatant filtration we used a 450-nm pore-size vacuum-driven Stericup filter (PVDF membrane) (Millipore, 20224462). For the concentration of supernatants, we used a 70 mL Centricon centrifugal filter (regenerated cellulose) with 100 kDa cut-off (Millipore, UFC710008). Gravity flow columns for the isolation of 2Strep-tagged vesicles included Strep-TactinXT 4Flow 1 mL (2-5012-001) and Strep-TactinXT 4Flow 5 mL (2-5013-001), along with Strep-TactinXT Buffer Set (2-1043-000; all from IBA Lifesciences). For the concentration and buffer exchange of isolated EVs, 15 mL Amicon Ultra-15 centrifugal filter units (regenerated cellulose) with 10 kDa cut-off were used (Millipore, UFC901008).

#### Material for precipitation-based extracellular vesicle isolation

A commercial kit for total exosome isolation from cell culture medium was used (Invitrogen, 4478359).

## Methods

### Cell culture

Cells used in this study were cultured in Dulbecco’s Modified Eagle Medium (DMEM, Thermo Fisher Scientific), supplemented with 2 mM L-glutamine, non-essential amino acids, 100 U/mL penicillin, 100 µg/mL streptomycin, and 10% fetal calf serum (FCS) (referred to as DMEMcplt). All cells were maintained in a humidified incubator at 37°C with 5% CO_2_.

EV-depleted FCS was prepared by ultracentrifugation of regular FCS at 120,000 *x g* for 18 hours using a swinging bucket rotor (SW40 Ti, Beckman Coulter). Approximately 10 mL of serum from the top layer of each centrifuge tube (Beckman Coulter, 344060) was carefully collected to avoid disturbing the pelleted EVs. This FCS was used to prepare EV-depleted DMEMcplt.

### DNA plasmid constructs

The pWPI-CD63-Nluc construct with the Blasticidin resistance gene (BLR) was generated by overlap extension PCR. Briefly, the Addgene plasmid CD63-pEGFP C2 (plasmid #62964) was used as template to amplify the CD63 coding region. This amplicon was fused at its 3’ end with the Nluc coding sequence derived from the Addgene plasmid pTRE3G-NlucP (plasmid #162595) via a short linker sequence (gagttc). The fused CD63-Nluc coding sequence was inserted into the lentiviral vector pWPI_ApoE-mTurquoise2 [2] via the BamHI and SpeI restriction sites. The resulting vector pWPI-CD63-Nluc (BLR) was used as a template for generating constructs encoding CD63 with internal insertion of the 2Strep-tag by using overlap extension PCR. Primers encoding the 2Strep-tag (underlined sequences) and flanking linkers: (SSGSA<u>WSHPQFEK-</u>GGGSGGGSGGSA-<u>WSHPQFEK-</u>GGGSGGGSGGSA) were used and amplicons were inserted into the large extracellular loop (LEL) of CD63 after serine residues 159 or 161, or after glycine residue 176, or into the small extracellular loop (SEL) after glutamine residue 36. The pWPI-CD63-Nluc construct with the 2Strep-tag after serine residue 159 is referred to as pWPI-CD63-2Strep-Nluc.

The plasmid pWPI-CD63-2Strep-PhoCl_2_-mScarlet-NLS, a lentiviral expression construct in which the CD63-2Strep coding region was fused at the 3’ end with the coding sequences of the photocleavable protein PhoCl_2_, mScarlet, and a nuclear localization signal (NLS) was generated by overlap extension PCR. As templates, we used the Addgene plasmid #164051 for the PhoCl_2_ coding sequence, the vector pWPI-ApoE-mScarletC1 for mScarlet [2], and a synthetic oligonucleotide coding for an NLS (PKKKRKVPKKEKG). The fused sequence was used to replace the Nluc coding region in pWPI-CD63-2Strep-Nluc.

The pWPI-CD63-Nluc-2Strep construct (BLR), in which the Nluc-2Strep sequence is fused to the C-terminus of wild-type CD63, and the pWPI-2Strep-Nluc-CD63 construct (BLR), in which the 2Strep–Nluc sequence is fused to the N-terminus of CD63, were generated using overlap extension PCR.

The pWPI-CD9-Nluc construct (BLR) was created by amplifying the CD9 coding sequence from Addgene plasmid CD9-mEGFP (plasmid #182864) and inserting it into the pWPI vector. The pWPI-CD9-2Strep-Nluc (BLR) construct was generated by inserting the 2Strep-tag with linkers after serine residue 164 of CD9.

### Generation of cell lines

Stable cell lines expressing engineered CD63 constructs with the 2Strep-tag, as well as control cells, were generated by lentiviral transduction. For production of lentivirus stocks, HEK293T cells were seeded into 10 cm-diameter dishes at a density of 5.5 x 10^6^ cells per dish in 5 mL DMEMcplt and on the next day, the medium was replaced with fresh DMEMcplt. For transfection, 8.4 µg of the lentiviral vector pWPI, 8.4 µg of the HIV-Gag packaging plasmid pCMV-dR8.91, and 4.2 µg of the vesicular stomatitis virus G glycoprotein (VSV-G) encoding plasmid pMD2.G were diluted in 700 µL OPTI-MEM (tube 1). In parallel, 44.5 µL of PEI (Polyethylenimine, 1 mg/mL; Polysciences Inc.) was mixed with 700 µL OPTI-MEM (tube 2). The contents of tube 1 and tube 2 were combined and incubated for 15 minutes before being added dropwise to the cells. Lentivirus-containing supernatants were collected approximately 48 hours post-transfection and filtered through a 0.45 μm pore-size filter (MF-Millipore). For cell line generation, Huh7 cells (1 x 10^5^ cells per well) and HEK293T cells (2 x 10^5^ cells per well) were plated in a 6-well plate and incubated with ∼1 mL of lentivirus-containing supernatant. At 24 hours post-inoculation, transduced cells were selected with 30 µg/mL Blasticidin (Gibco, 46-1120) for a minimum of three weeks to ensure stable expression of the transgene. To maintain stable expression, cells were cultured in medium containing 10 µg/mL Blasticidin.

### Preprocessing of supernatants for extracellular vesicles isolation

Four confluent 15 cm-diameter dishes of Huh7 cells stably expressing CD63-2Strep-Nluc, or control cells, were trypsinized, washed twice with PBS, pelleted, and resuspended in 30 mL of EV-depleted DMEMcplt. For each condition, cells were seeded at a density of 2.5 x 10^6^ cells per 15 cm-diameter dish in 15 mL of EV-depleted DMEMcplt (10 dishes per condition). After 72 hours, supernatants were collected and ∼110 mL of supernatant was filtered using a 450 nm pore size Stericup PVDF filter. Two 55-mL portions of the filtered supernatants were further concentrated to ∼2 mL and free proteins were removed by centrifugation using a Centricon 70 (100 kDa cut-off) at 3,500 *x g* at 4°C for a total of ∼65 minutes. Concentrated supernatant was diluted in 1x Buffer W (100 mM Tris-HCl, pH 8.0, 150 mM NaCl, 1 mM EDTA) to a final volume of 12.5 mL (input supernatant) that was used for EV isolation.

### Affinity-based isolation of extracellular vesicles

For affinity-based isolation of EVs produced in Huh7-derived cells, 8 mL of input supernatant was added to a 5 mL gravity-flow column packed with Strep-TactinXT resin, which was further processed according to the manufacturer’s protocol with slight modifications. In brief, the Strep-TactinXT 4Flow column was equilibrated with a two-column volume of 1x Buffer W. The input supernatant was applied to the column and allowed to flow through by gravity. The column was washed with 4 x 2 column volumes of 1x Buffer W, and elution was carried in four sequential steps, each using 0.8 column volumes of 1x Buffer BXT (100 mM Tris-HCl, pH 8.0, 150 mM NaCl, 1 mM EDTA, 50 mM biotin), with the eluate collected in 0.8 CV fractions. The four eluate fractions (16 mL total) were pooled, and 10 mL of the pooled eluate was concentrated using an Amicon Ultra-15 filter with a 10 kDa cut-off at 3500 *x g*, reducing the volume to ∼640 µL. All fractions, including input, flow-through, washes, and eluates, were collected for nanoluciferase (Nluc) measurement to monitor the isolation process.

For regeneration of the column, it was washed with 8 column volumes of Buffer XT-R (3 M MgCl_2_). To assess column regeneration, 5 mL of 1x HABA containing solution (100 mM Tris-HCl, pH 8.0, 150 mM NaCl, 1 mM EDTA, 1 mM HABA) (IBA Lifesciences, catalogue number 2-1002-100) was added, with a yellow-to-orange color shift indicating successful regeneration. Right thereafter, the column was washed with 8 column volumes of 1x Buffer W and overlaid with 2 mL of 1x Buffer W. Finally, the column was sealed with caps, wrapped in parafilm, and stored at 4°C.

Affinity-based isolation of EVs produced in HEK293T-derived cells was performed using the protocol described above, but cells were seeded at a density of 5-6 x 10^6^ cells per 15-cm diameter dish, and the entire input supernatant sample was used.

### Isolation of extracellular vesicles by the precipitation method

To 4 mL of input supernatant, 4 mL of total EV isolation reagent (Thermo Fisher Scientific) was added and the mixture was incubated overnight at 4°C. On the next day, the samples were centrifuged at 10,000 *x g* for 60 minutes at 4°C. The supernatant was carefully aspirated and discarded, while total EVs retained in the pellet were resuspended in 512 µL of 1x Buffer W to match the input volume processed in the affinity-based method. For resuspension, the sample was gently vortexed for ∼5 minutes.

### Specificity index of affinity-based isolation of extracellular vesicles

To calculate the specificity with which engineered EVs can be isolated by the 2Strep-based method, we defined a specificity index (SI). The SI was determined using CD63-2Strep-Nluc EVs and non-2Strep-tagged EVs (CD63-Nluc) as follows:

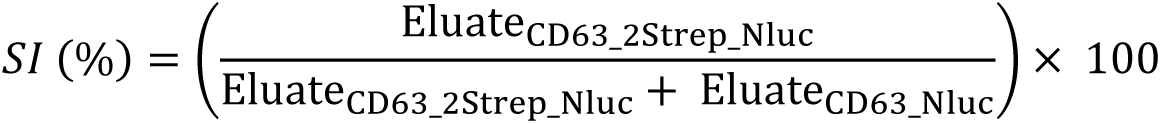

### Production and isolation of VSV-G coated extracellular vesicles

HEK293T cells stably expressing CD63-2Strep-mScarlet (Fig. 1B) were seeded onto 10 15-cm diameter dishes at a density of 5 x 10⁶ cells per dish in DMEMcplt. At ∼40 hours post-seeding, cells were transiently transfected with 30 µg of the plasmid pMD2.G encoding VSV-G using the PEI transfection reagent (DNA:PEI ratio 1:3). After 24 hours, the medium was replaced with 15 ml of EV-depleted DMEMcplt, and cells were cultured for an additional 48 hours. Cell-conditioned medium was collected, preprocessed as described above, and subjected to affinity-based isolation using the entire input supernatant. EVs were subsequently concentrated and the buffer was exchanged against 1x PBS using Amicon Ultra-15 filter with a 10 kDa cut-off. To detach the mScarlet-NLS from the CD63 scaffold, EVs were illuminated with 405 nm light using a 6W UV LED lamp (Sovol, 358 mW/cm², model GHD-XZBDS) under continuous rotation for 5 minutes to induce PhoCl_2_-mediated cleavage.

### Luciferase reporter assay

To measure Nanoluciferase (Nluc) activity in CD63-Nluc-loaded EVs, samples were diluted ∼5 times using 1x buffer W (IBA Lifesciences) and then mixed with an equal volume of 0.1% Triton X-100 in PBS for permeabilization, resulting in a final working concentration of 0.05% Triton X-100 to enhance substrate penetration. Diluted samples (50 µL) were transferred to black glass-bottom 96-well plates. An equal volume of Nluc substrate (Nano-Glo Luciferase Assay System, Promega), diluted 1,000-times in luciferase assay buffer (25 mM glycylglycine, 15 mM MgSO_4_, 4 mM EGTA, 1 mM DTT, 15 mM K_3_PO_4_, pH 7.8), was added to each well. Nluc activity was measured using a Mithras LB 940 plate luminometer (Berthold Technologies, Freiburg, Germany) using the following setting: 10-second shake and 1-second signal counting.

### Hepatitis C virus infection and viral replication quantification

Huh7 cells stably expressing CD63-2Strep-Nluc, or CD63-Nluc control cells, were seeded into 6-well plates at a density of 1 x 10^5^ cells per well (2 wells per conditions). On the next day, cells were inoculated with 2 mL of HCV (strain Jc1; titer ∼2 x 10^6^ Tissue Culture Infectious Dose 50% (TCID_50_)/mL determined on Huh7.5 cells as described earlier [55]). At 4 hours post-infection, the medium was replaced with 2 mL of fresh DMEMcplt per well. At 72 hours post-infection, cell culture supernatants were collected and 1/10 volume of a 10x stock of Buffer W was added to adjust the pH value. A total of 3.2 mL of this supernatant was used for capturing of CD63-2Strep-tagged EVs encapsulating Nluc using a 1 mL Strep-TactinXT column. The column was washed with 6 column volumes of 1x Buffer W and eluted with 3.2 column volumes of 1x Buffer BXT (100 mM Tris-HCl, pH 8.0, 150 mM NaCl, 1 mM EDTA, 50 mM biotin). All fractions, including input, flow-through, wash, and eluate were treated with an equal volume of 1% Triton X-100 in PBS to inactivate viral infectivity prior to Nluc activity measurement and HCV core protein quantification. To quantify core protein, we used a chemiluminescent microparticle immunoassay (CMIA) with the ARCHITECT HCV Ag Reagent Kit (6L47; Abbott Diagnostics) according to the manufacturer’s instructions.

### Live-cell spinning-disk confocal microscopy

Huh7 cells stably expressing CD63-2Strep-mScarlet were seeded onto 35-mm diameter glass-bottom imaging dishes (MatTek Corporation). Prior to imaging, cells were washed twice with PBS and either mock-treated or exposed to 80 µM Colchicine in phenol red-free DMEMcplt, supplemented with 25 mM HEPES (pH 7.4). Time-lapse live-cell imaging was performed in a humidified chamber at 37°C and 5% CO_2_ using the PerkinElmer spinning-disk confocal microscope system with an Apo TIRF 60x/1.49 N.A. oil immersion objective, as previously described [2]. A solid-state laser with an excitation at 561 nm and corresponding emission filter was employed. Imaging was performed at 0.56-second intervals for a total duration of 1 minute.

### Uptake analysis of isolated 2Strep-tagged extracellular vesicles encapsulating reporter proteins

A pooled eluate containing 2Strep-tagged EVs that were isolated from cells stably expressing CD63-2Strep-mScarlet or CD63-2Strep-Nluc by the Strep-based method, was concentrated to ∼500 µL using an Amicon Ultra-15 filter with a 10 kDa cut-off at 3,500 *x g*, and the buffer was exchanged using 15 mL of 1x PBS. Total EVs isolated via the precipitation method from the same cells or from naïve cells, were processed in parallel.

Uptake analysis of Huh7-derived mScarlet-containing EVs was performed using Huh7 cells expressing eYFP-CaaX that labels the plasma membrane and endosomes [2]. Briefly, 1,400 cells in 50 µL of DMEMcplt were seeded into a µ-slide 15-well chambered coverslip (Ibidi, catalogue number 81506). At 24 hours post-seeding, the medium was replaced with 40 µL of phenol red-free DMEMcplt, and cells were treated with 15 µL of 2Strep-EVs (total protein amount not quantifiable because it was below the linear detection range of the Bradford assay) or control EVs isolated by the precipitation method from the culture supernatant of naïve Huh7 cells (33 µg). After 6 hours of incubation, cells were imaged using live-cell spinning disk confocal microscopy for 1 minute at 1.43-second interval. A solid-state laser with excitation at 488 nm and 561 nm and corresponding emission filter were employed. The first frames of the acquired images were used to quantify the number of fluorescent foci taken up by the cells after background subtraction, using the ColocQuant software [56] with a threshold of 5.

Dose-dependent uptake analysis of HEK293-derived 2Strep-EVs containing mScarlet was performed the same way with minor modifications. Briefly, cells were treated with 22 µL of 2Strep-EVs (corresponding to 2.5 x 10^5^ EVs per cell) or with serially diluted EV samples as specified in Fig. S2E. After 24 hours of incubation, live-cell imaging was performed, capturing a 2 µm z-stack with 0.2 µm step intervals. Maximum intensity projections were used to quantify internalized mScarlet signal based on mean fluorescence intensity. For western blot analysis, cells were seeded into 96-well plates at a density of 1.5 x 10^4^ cells per well and treated with various amounts of EVs, starting at 2.5 x 10^4^ EVs per cell as specified in Fig. 6G. After 24 hours, cells were harvested and analyzed by Western blot to detect internalized EVs using CD63- and RFP-specific antibodies.

For uptake analysis of Huh7-derived 2Strep-EVs containing Nluc, 1 x 10^4^ Huh7 cells were seeded into each well of a white 96-well plate (Greiner bio-one, 655098) in 100 µL DMEMcplt. At 24 hours post-seeding, cells were incubated with 20 µL of affinity-isolated 2Strep-EVs (total protein amount not quantifiable) or EVs (19.4 µg) isolated with the precipitation method at 37°C for 2 or 6 hours, or at 4°C for 2 hours (using plates pre-cooled on ice for 30 minutes). Thereafter, cells were washed four times with PBS and lysed in 100 µL PBS containing 0.5% Triton X-100. Cell-conditioned media were also collected and adjusted to 0.5% Triton X-100 in PBS to assess the amount of non-internalized 2Strep-EVs. The percentage of EVs taken up into cells was calculated by determining the ratio of intracellular to the total Nluc activity (intra- and extracellular).

### Immunofluorescence and confocal microscopy

Immunofluorescence was performed as previously described [56]. All steps were conducted at RT and cells were washed at least three times with PBS between each incubation step. Briefly, cells cultured on glass coverslips were fixed in 4% paraformaldehyde (PFA) in PBS for 10 minutes. Permeabilization was carried out using 0.1% Triton X-100 in PBS for 10 minutes. Cells were incubated with 3% (w/v) bovine serum albumin (BSA) in PBS for 20 minutes and then with primary antibodies in 1% BSA/PBS for 1 hour (dilution of rabbit anti-LAMP1 1:250 and mouse anti-NS5A 1:500). Secondary antibodies conjugated to Alexa Fluor 488 or 647 (diluted 1:1000) were applied for 1 hour. For the detection of endogenous CD63, cells were stained with an anti-CD63 antibody conjugated to Alexa Fluro 488 (1:400). Nuclei were visualized with DAPI staining (1:1000; Molecular Probes). After staining, coverslips were mounted onto glass slides with Fluoromount-G mounting medium (Invitrogen, catalogue number 00-4958-02) and left overnight at 4°C before imaging. Images were acquired with a spinning-disk confocal microscope (PerkinElmer).

The fluorescence intensity profiles for each channel were obtained using the “Plot Profile” function in Fiji [57]. The raw intensity values of the selected reference channel were kept unchanged, while the intensity values for the second channel along the same line were normalized by adjusting them based on the ratio of the mean intensity of the reference channel to that of the second channel.

### Western blot analysis

Isolated EVs were lysed in SDS-containing sample buffer (final concentration: 62.5 mM Tris-HCl, pH 6.8, 0.02% Bromophenol Blue, 3.3% glycerol, 0.5% SDS and 0.33% β-mercaptoethanol). Cell lysates were prepared in 2x sample buffer containing 5 mM MgCl₂ and 5 U/mL of benzonase, and incubated at 37°C for 5 minutes. Samples were further denatured by 5 minutes treatment at 95°C. Proteins were separated by SDS-PAGE and transferred to a polyvinylidene difluoride (PVDF) membrane that was incubated in 5% skim milk in PBS-0.05% Tween 20 (PBST), pH 7.4, for one hour at RT. Subsequently, the membrane was incubated with primary antibody in 1% skim milk-containing PBST overnight at 4°C, prior to incubation with a secondary antibody conjugated to horseradish peroxidase (HRP) for one hour at RT. Bound secondary antibodies were detected using the Western Lightning Plus-ECL system (PerkinElmer), and the emitted chemiluminescent signals were captured with an Intas ChemoCam Imager 3.2 (Intas).

### Negative staining and transmission electron microscopy (TEM)

Pioloform-coated 300-mesh copper grids (Science Services GmbH, Munich, Germany) were coated with a 2.7 nm-thick carbon layer using a Leica ACE600 carbon coater to enhance the stability of the support film. Thereafter, the grids were freshly glow-discharged and placed on top of 5-7 µL drops of isolated EVs for 5 minutes at RT to allow for sample absorption. The grids were quickly washed with water, followed by a brief dip and a 5-minute incubation on top of drops of 3% uranyl acetate in water at RT. After staining, excess liquid was gently removed by blotting with filter paper and the grids were air-dried. Imaging was performed on a Jeol JEM-1400 transmission electron microscope (Jeol Ltd., Tokyo, Japan), equipped with a 4k pixel digital camera (TemCam F416; TVIPS, Gauting, Germany).

### Size determination of extracellular vesicles by transmission electron microscopy

Electron micrographs of EVs visualized by negative staining and TEM were segmented using the Ilastik software [58] with the Autocontext workflow involving two stages of pixel classification. The resulting binary masks were analyzed in Fiji to measure EV diameters that were calculated as the average of the major and minor axes of fitted ellipses.

### Nanoparticle tracking analysis (NTA)

For quantification and size distribution of EVs, samples were diluted 10-20 times in 1x PBS and analyzed using on a Nanosight NS300 (Malvern Panalytical, Malvern, UK). For each sample, 3 separate runs of 30 seconds were recorded. For the analysis of Brownian motion of particles in solution, Nanosight NTA software v3.3 at a camera level of 9 and detection threshold of 2 or 3 was used. Size and concentration data from each recorded run were averaged to yield values representative for each sample. Values were corrected for individual dilution.

### Quantification of CD63 protein amount by ELISA

The amount of CD63 protein in mScarlet-EVs produced in HEK293T cells was quantified using the Human CD63 ELISA kit (RayBiotech, ELH-CD63-2-RB) according to the manufacturer’s protocol. Standards were prepared ranging from 12,500 pg/ml to 390.6 pg/ml using 2-fold serial dilutions. The EV samples were diluted to fall within the linear detection range of the standards.

### Correlative light and electron microscopy (CLEM)

Huh7 cells stably expressing CD63-mScarlet-NLS were analyzed by CLEM using lipid droplets stained with LipidTOX (ThermoFisher Scientific, catalogue number H34477) as fiducial markers, as previously described [2]. Large lipid droplets (>500 nm) with distinct landmarks were used for low precision correlation of IF and EM micrographs using the Landmark Correspondences plugin in the Fiji software package. For high-precision correlation, smaller lipid droplets (≤200 nm) appearing as spherical structures in no more than three consecutive 70 nm-thick EM sections were used.

### Flotation of isolated extracellular vesicles using iodixanol density gradient centrifugation

To prepare the iodixanol gradient, OptiPrep (60% iodixanol in water (w/v), Sigma-Aldrich, catalogue number D1556) was sequentially diluted in PBS to obtain final concentrations of 50%, 40%, 30%, 20%, and 10%. A volume of 700 µL of 50% iodixanol was placed at the bottom of the centrifugation tube (Beckman coulter, catalogue number 344062), frozen at −80°C, and subsequently overlaid with 700 µL of each lighter fraction, with freezing between each step to stabilize the gradient. For flotation, 150 µL of isolated EVs were mixed with 750 µL of OptiPrep to reach a final concentration of 50% iodixanol. This sample was placed at the bottom of the thawed gradient using a 2 mL syringe fitted with a 70 mm, 20-gauge needle. The samples were centrifuged at 208,000 *x g* for 16 hours at 4°C using an SW60 rotor (Beckman Coulter, Inc.) with an acceleration and deceleration rate of 5. Eleven fractions were collected from top to bottom, diluted twofold with 0.1% Triton X-100 in PBS, and analyzed for Nluc activity. The refractive index of each fraction was measured using a refractometer (Kruess, AGS Scientific, Germany).

### Proteomic analysis of isolated 2Strep-tagged extracellular vesicles

*Sample preparation:* 10 µL were used for the EV samples prepared by the precipitation method (Prec-EV), and 100 µL for the 2Strep-tag-isolated EV samples (2Strep-EV). A lower volume was used for Prec-EVs due to their high protein content, which would otherwise overload the mass spectrometry system and compromise detection sensitivity. Both samples were dried down in a SpeedVac and resuspended in 20 µL of RIPA buffer containing 2% SDS. Samples were digested with trypsin using an AssayMAP Bravo liquid handling system (Agilent technologies) running the autoSP3 protocol according to Müller and colleagues [59].

*MS method Ultimate 3000 coupled to an Orbitrap Exploris 480 equipped with a FAIMS Pro:* the dried peptide sample was reconstituted in 97.4% water, 2.5% hexafluoro-2-propanol and 0.1% trifluoroacetic acid and one third of the sample was used for the analysis. The LC-MS/MS analysis was carried out on an Ultimate 3000 UPLC system (Thermo Fisher Scientific) directly connected to an Orbitrap Exploris 480 mass spectrometer. The instrument operated in positive mode with the FAIMS Pro interface, applying a compensation voltage of −45 V to exclude singly charged ions. The inner and outer electrode temperatures were set to default, and nitrogen was used as the carrier gas at the default flow rate. Each sample was analyzed for a total of 60 min. Peptides were online desalted on a trapping cartridge (Acclaim PepMap300 C18, 5µm, 300Å wide pore; Thermo Fisher Scientific) for 3 min using 30 µL/min flow of 0.05% TFA in water. The analytical multistep gradient (300 nL/min) was performed using a nanoEase MZ Peptide analytical column (300Å, 1.7 µm, 75 µm x 200 mm, Waters) using solvent A (0.1% formic acid in water) and solvent B (0.1% formic acid in acetonitrile). For 45 min the concentration of solvent B was linearly ramped from 4% to 30%, followed by a quick ramp to 78%; after two minutes the concentration of solvent B was lowered to 2% and a 10 min equilibration step appended.

*Data-independent acquisition (DIA)*: eluting peptides were analyzed in the mass spectrometer using DIA mode. A full scan at 120k resolution (380-1400 m/z, 300% AGC target, 45 ms maxIT) was followed by 20 DIA windows. The DIA acquisition covered a mass range of 400-1000 m/z using windows of a variable width. Windows overlapped by 1 m/z, the AGC target was 1000% with a maxIT of “auto” and spectra were recorded at a resolution of 60k. After each sample, a wash run (20 min) was performed to minimize carry-over between samples. Throughout the course of the measurement, instrument performance was monitored by regular (∼one per 48 hours) injections of a standard sample and an in-house shiny application.

*Data analysis:* the DIA RAW files were analyzed with Spectronaut (Biognosys, version 19.9.250422.62635) [60] in directDIA+ (deep) library-free mode. Default settings were applied with the following adaptions. Within DIA analysis under identification, the Precursor PEP Cutoff was set to 0.01, the Protein Qvalue Cutoff (Run) set to 0.01 and the Protein PEP Cutoff set to 0.01. In Quantification, the Proteotypicity Filter was set to Only Protein Group Specific, the Protein LFQ Method was set to MaxLFQ, Cross-Run Normalization was set to False and the Quantification window was set to Not Synchronized (SN 17). In Workflow, Method Evaluation was set to True. The data was searched against the human and *bos taurus* proteome from Uniprot (human and *bos taurus* reference databases with one protein sequence per gene, containing 20,597 unique entries from 21^st^ of January 2025 and 26,942 unique entries from 24^th^ of March 2025, respectively) and the contaminants FASTA from MaxQuant (246 unique entries from 22^nd^ of December 2022).

Conditions were included in the setup. The mass spectrometry proteomics data have been deposited to the ProteomeXchange Consortium via the PRIDE [61] partner repository with the dataset identifier PXD071011 and the token HurQZjbYjkpP.

### Statistics

For the statistical analysis log_2_ transformed AvgIBAQ quantity-values were used. Protein groups retaining valid values in at least 70% of the samples in one or more conditions were included. No across-sample normalization was applied. Adapted from the Perseus recommendations [62] missing values being completely absent in one condition, were imputed with random values drawn from a downshifted (2.2 standard deviation) and narrowed (0.3 standard deviation) intensity distribution of the individual samples. For partially missing values (not completely absent in any condition), imputation was performed using the R package missForest [63]. Differential abundance was analyzed using the R-package “limma” [64] with eBayes options ‘trend’ and ‘robust’ to TRUE, adapting the pipeline for proteomics data. Due to violation of homoskedasticity in the EV data, statistical testing was conducted using a protein pairwise contrast, in which all possible between conditions protein differences were compared in in a paired (within sample) manner. The p-values were adjusted with the Benjamini–Hochberg method for the multiple testing [65]. For each protein the median log fold change and median adjusted p-value, from all pairwise comparisons were calculated and reported. This approach obviates the need for external normalization.

### Motility analysis of CD63-2Strep-mScarlet

Tracking of CD63-2Strep-mScarlet-NLS in fluorescence microscopy image data was performed by probabilistic particle tracking [66]. The method is based on Kalman filtering and particle filtering, and integrates multiple measurements by separate sensor models. Elliptical sampling [67] was used to obtain detection-based and prediction-based measurements, which were integrated by a sequential multi-sensor data fusion method that takes separate uncertainties into account. Motion information was exploited by a displacement term in the cost function, and information from both past and future time points was combined using Bayesian smoothing. Particles were detected using the spot-enhancing filter (SEF) [68], which involves applying a Laplacian of Gaussian (LoG) filter to the fluorescence microscopy images, followed by thresholding the filter response and determining the connected components. The threshold was automatically computed as the mean of the filter response plus two times the standard deviation.

To quantify the motility of the tracked particles, a mean-square displacement (MSD) analysis [69] was performed. The MSD values were computed as a function of the time intervals Δt for each trajectory with a minimum time duration of about 2.2 s (4 time steps). To obtain MSD curves for each experimental condition, all corresponding MSD curves were averaged.

### Statistical analysis

Unless otherwise stated, statistical comparisons between sample groups were performed using a two-tailed Mann-Whitney test in GraphPad Prism 8. Differences with p-values less than 0.05 were considered statistically significant and are indicated in the graphs.

## Acknowledgments

We acknowledge Ulrike Herian and Marie Bartenschlager for their technical assistance; the University of Heidelberg Electron Microscopy Core Facility (EMCF, Heidelberg, Germany), co-headed by Dr. Charlotta Funaya and Dr. Réza Shahidi, for their continuous support; and the Infectious Diseases Imaging Platform (IDIP), headed by Dr. Vibor Laketa at the Center for Integrative Infectious Disease Research (CIID, Heidelberg, Germany). We are grateful to Dr. C.M. Rice for the monoclonal antibody 9E10 and acknowledge the following persons for the provision of plasmids via Addgene: CD63-pEGFP C2 (#62964), a gift from Paul Luzio; pTRE3G-NlucP (#162595) from Masaharu Somiya; plasmid pcDNA-NBid-PhoCl_2_-CBid (#164051) from Robert Campbell, and plasmid CD9-mEGFP (#182864) from Salvatore Chiantia. Lastly, we sincerely appreciate the insightful discussions and support from all members of the Molecular Virology unit.

## Data availability statement

The datasets generated from the proteomics analysis of isolated EVs are available in the PRIDE repository under the project accession number PXD071011 with the token HurQZjbYjkpP.

## Funding information

This work was supported by grants from the Deutsche Forschungsgemeinschaft (DFG, German Research Foundation) – Project Number 272983813 – TRR 179 to RB and Project Number 240245660 - SFB1129 to RB and KR as well as Project Number 529989072 to KR. The work was also supported by the German Center for Infection Research (DZIF), project number DZIF-TTU Imag 5.712 and TTU05.821, both to RB. The funders had no role in study design, data collection and analysis, decision to publish, or preparation of the manuscript.

## Conflict of interest disclosure

The authors declare no conflicts of interest.

## Supplementary information

**Figure S1.**
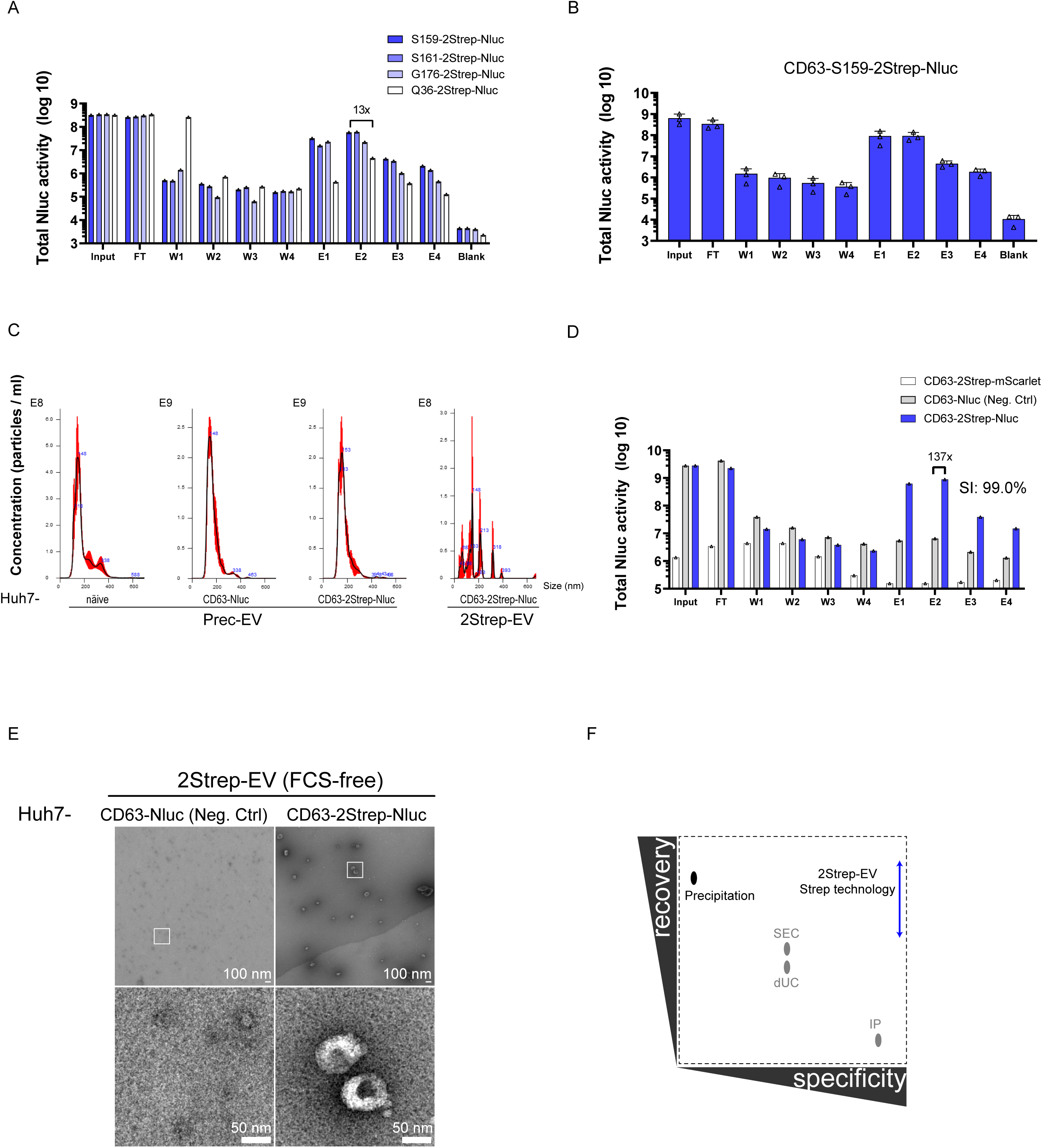
**Identification of an internal CD63 tagging site and characterization of twin-Strep tagged EVs** (A) Screening for functional 2Strep-tag insertion sites within CD63. Huh7 cells stably expressing C-terminally Nluc-tagged CD63 with 2Strep-tag insertions at given sites in the LEL (S159, S161, G176, or Q36) were generated by lentiviral transduction. Culture supernatants from these cells were used to isolate EVs by the Strep-based method. Nluc activity was measured in all fractions collected during the isolation process (flow-through, FT; washes, W; eluates, E). Values were calculated for the total volume of each fraction and are displayed on a logarithmic scale. “Blank” represents assay background determined using PBS. N=1. The same results were obtained in pilot experiments using Strep-TactinXT magnetic beads (not shown). (B) Confirmation of the suitability of CD63-Nluc with the 2Strep insertion at position S159. Data represent means (SD) from three independent biological experiments. (C) Nanoparticle tracking analysis of EVs isolated with the precipitation method (Prec-EVs) or the 2Strep-based approach (2Strep-EVs) as described in Fig. 3. Shown are the size distribution and particle concentration of EVs containing the CD63 tagged at position S159. Data are displayed as means (SEM, in red) from 3 separate runs for each sample. (D) Isolation of EVs from the supernatant of cells cultured in FCS-free medium. Huh7 cells stably expressing the indicated CD63-2Strep proteins were cultured in FCS-free media for 24 hours. The cell-conditioned media were membrane-filtered (450 nm) and concentrated using Centricon spin columns (100 kDa cut-off) prior to 2Strep affinity-based isolation. Nluc activity was measured in all fractions including input, flow-through (FT), washes (W), and eluates (E) to assess the efficiency of the EV isolation process. Values were calculated for the total volume and are presented on a logarithmic scale. Neg. Ctrl, negative control corresponding to a non-2Strep-tagged CD63. SI: specificity index. (E) Isolated EVs from (D) were visualized by negative staining and TEM. The boxed area in the overview electron micrograph (upper panel) is magnified in the lower panel, showing highly enriched EVs of small sizes. (F) Schematic illustrating EV recovery versus specificity. Using the 2Strep technology, 2Strep-tagged CD63-containing EVs could be enriched with ∼40-80% recovery and ∼97% specificity. Recovery and specificity values achieved by using differential ultracentrifugation (dUC), size exclusion chromatography (SEC), and immunoprecipitation are shown for comparison and are displayed according to the MISEV guidelines [3].

**Figure S2.**
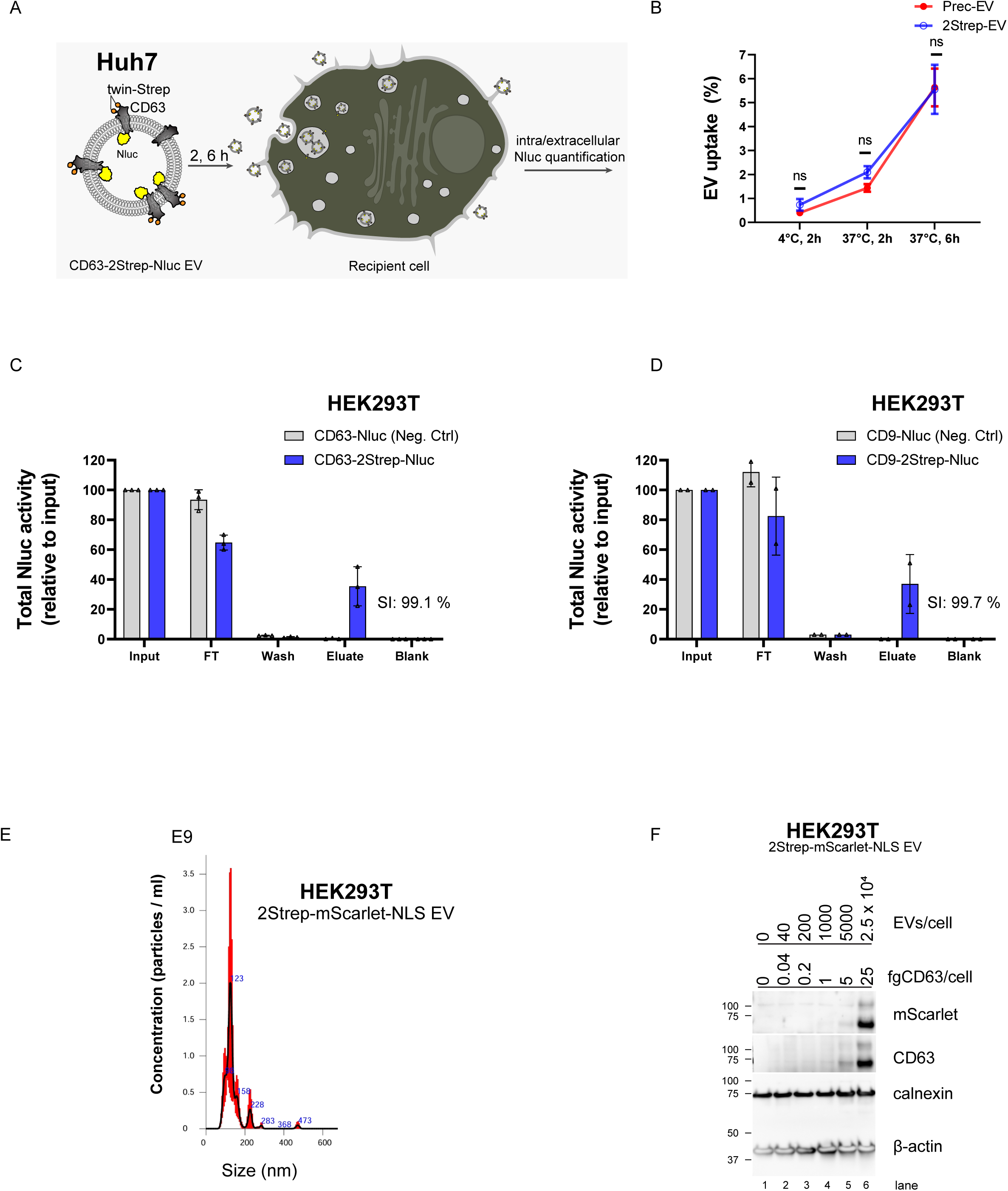
**Cellular uptake of affinity-isolated 2Strep-EVs containing reporter proteins** (A) Schematic depicting the experimental procedure. Nluc-equivalent amounts of Huh7-derived EVs encapsulating Nluc isolated by precipitation (Prec-EVs) or the Strep-based method (2Strep-EVs) (refers to Fig. 3) were added to Huh7 cells for 2 hours at 4°C (adsorption to the cells) or 37°C, and for 6 hours at 37°C. Thereafter, Nluc activity was measured in cell lysates and culture supernatants (intracellular and extracellular Nluc, respectively). (B) Quantification of EV uptake. Values (%) refer to the proportion of intracellular Nluc activity relative to the sum of intracellular and extracellular Nluc activity. Data represent means (SD) from three independent biological experiments. Statistical significance was determined using a two-tailed Mann-Whitney test. *(C-F) Isolation and dose-dependent uptake analysis of HEK293T-derived 2Strep-EVs containing reporter proteins into Huh7 cells* (C) Isolation of CD63-2Strep-Nluc EVs from HEK293T cell culture supernatant. Culture supernatants (5 mL) from HEK293T cells stably expressing CD63-2Strep-Nluc or CD63-Nluc control cells (Neg. Ctrl) were subjected to Strep-TactinXT affinity purification. Nluc activity was measured in all fractions collected during the isolation process (flow-through, FT; washes, W; eluates, E). Values were normalized to the total volume and are shown relative to the input that was set to 100%. Data represent means (SD) from three independent biological experiments. SI: specificity index. (D) Isolation of CD9-2Strep-Nluc EVs from HEK293T cell culture supernatant. Culture supernatants (5 mL) from HEK293T cells stably expressing CD9-2Strep-Nluc or CD9-Nluc control cells (Neg. Ctrl) were subjected to Strep-TactinXT affinity purification. Nluc activity was measured in all fractions collected during the isolation process (flow-through, FT; washes, W; eluates, E). Values were normalized to the total volume and are shown relative to the input that was set to 100%. Data represent means (SD) from two independent biological experiments. (E) Nanoparticle tracking analysis of 2Strep-EVs encapsulating mScarlet-NLS, isolated from culture supernatant of HEK293T-derived cells. Data are displayed as the mean (SEM, in red) from 3 separate runs of the same sample. (F) Huh7-derived cells expressing the membrane marker eYFP-CaaX were incubated with increasing amounts of EVs (ranging from 0 to 2.5 x 10⁴ EVs per cell according to NTA or given amounts of CD63 in femtograms per cell) for 24 hours. EV uptake was analyzed by Western blot using RFP (mScarlet)- and CD63-specific antibodies. Calnexin and β-actin served as loading controls. fg, femtogram.

**Video 1. Intracellular trafficking of CD63-2Strep-mScarlet in mock versus colchicine-treated hepatocytes.**

Huh7 cells stably expressing CD63-2Strep-mScarlet were either mock-treated or treated with 80 µM colchicine for 1 hour and analyzed by live-cell spinning-disk confocal microscopy. The motility of mScarlet is shown on the left and right for mock- and colchicine-treated cells, respectively. Images were acquired at 0.56-second intervals over a total duration of 1 minute. A representative duration of 10.64 seconds is shown.

**Video 2. Internalization of isolated mScarlet-loaded 2Strep-EVs**

Huh7 cells stably expressing eYFP-tagged CaaX were incubated with Huh7-derived 2Strep-EVs containing mScarlet. At 6 hours post-incubation, cells were washed and imaged using live-cell spinning-disk confocal microscopy to visualize internalized fluorescent signals. A representative region within the cell boundary showing internalized mScarlet associated with putative endosomes is presented. The time-lapse sequence spans 12.87 seconds with a frame interval of 1.43 seconds. CaaX refers to the farnesylation signal derived from the human HRAS protein which targets eYFP to intracellular membranes.

## Notes

### Competing Interest Statement

The authors have declared no competing interest.

### Summary of Updates

This version of the manuscript has been revised to update the Title, Abstract, Figure 1, and Main Text. Notably, the manuscript now includes CD9, in addition to CD63, as a scaffold for EV engineering.

